# Disrupted TRiC allostery and conformational equilibrium by LCA-linked mutations: structural and functional insights

**DOI:** 10.64898/2026.01.22.701072

**Authors:** Qiaoyu Zhao, Caixuan Liu, Qing Zhang, Zuyang Li, Yanxing Wang, Xuehai Zhou, Xingyan Ye, Shanshan Wang, Yanke Wang, Wanying Jiang, Qianqian Song, Yao Cong

## Abstract

The eukaryotic chaperonin TRiC/CCT is essential for proteostatis, yet the molecular basis of its subunit-specific pathologies remains poorly understood. Here, we elucidate the molecular mechanism of Leber Congenital Amaurosis (LCA), a severe hereditary retinal dystrophy arised from mutations in the CCT2 subunit of TRiC. By integrating cryo-electron microscopy, biochemistry, and proteomics, we demonstrate that LCA-associated mutations (T400P and R516H in CCT2) disrupt TRiC’s critical intra-molecular and intra-ring allosteric network and impair its functional cycle, drastically reducing the population of the folding-active closed state. Unexpectedly, we captured a fully folded endogenous α-tubulin within the mutant TRiC chamber, revealing a unique CCT8 C-terminal tail involved folding pathway, distinct from β-tubulin. Furthermore, cellular proteomics revealed that TRiC dysfunction causes a specific downregulation of essential SLC membrane transporters. We propose that the loss of these transporters is likely catastrophic for metabolically demanding tissues like the retina and developing embryo. Our work provides a direct mechanistic link between TRiC structural defects and LCA pathology, offering a new framework for understanding etiology of chaperonopathies.

## Introduction

The maintenance of proteostasis is fundamental to cellular health, and its failure is a hallmark of numerous human diseases. Central to this network is the eukaryotic chaperonin TRiC/CCT, an ATP-dependent molecular machine essential for the folding of approximately 10% of the cytosolic proteome^1^, including cytoskeletal proteins such as actin and tubulin^2,3^. TRiC comprises two stacked rings, each consisting of eight paralogous subunits (CCT1 to CCT8)^4,5^. Each subunit contains an apical (A) domain for substrate recognition, an intermediate (I) domain for for allosteric signaling, and an equatorial (E) domain that powers the complex through ATP hydrolysis^6-8^. TRiC function relies on a precisely coordinated, ATP-driven conformational cycle that transitions the complex between open (substrate-accepting) and closed (folding-active) conformations. In recent years, mutations in CCT subunits have been linked to a growing spectrum of disorders termed “TRiCopathies”, with clinical manifestations ranging from sensory neuropathy to severe developmental defects^9-15^. Previous structural analysis of the hereditary sensory neuropathy variant CCT5-H147R, reconstituted as a homo-oligomeric complex, revealed that a de novo intramolecular hydrogen bond between R147 and S428 alters the flexibility of the CCT5 equatorial domain^16^. However, the structural effects of the pathogenic TRiC mutations in the native hetero-hexadecameric chaperonin remain poorly understood.

Leber Congenital Amaurosis (LCA) is a devastating inherited retinal dystrophy, characterized by profound vision loss at birth^17^. LCA has been linked to compound heterozygous mutations (T400P and R516H) in the CCT2 subunit of TRiC^15^, resulting in a heterogeneous population of mutant complexes (three different TRiC complexes) in patients (Fig. S1A). The high evolutionary conservation of these residues (Fig. S1B), along with the embryonic lethality of the homozygous T400P mutation in mice^18^, underscores their physiological importance. Structurally, these residues occupy critical functional regions: T400 resides in H11 in I-domain, adjacent to the catalytic D392, which is critical for ATP hydrolysis; while R516 is situated on H15 in the C-terminal region of E-domain, at the inter-subunit interface (Fig. 1A). However, the molecular mechanism linking these specific point mutations to TRiC dysfunction and subsequent LCA retinal pathology remains unclear.

**Fig. 1.**
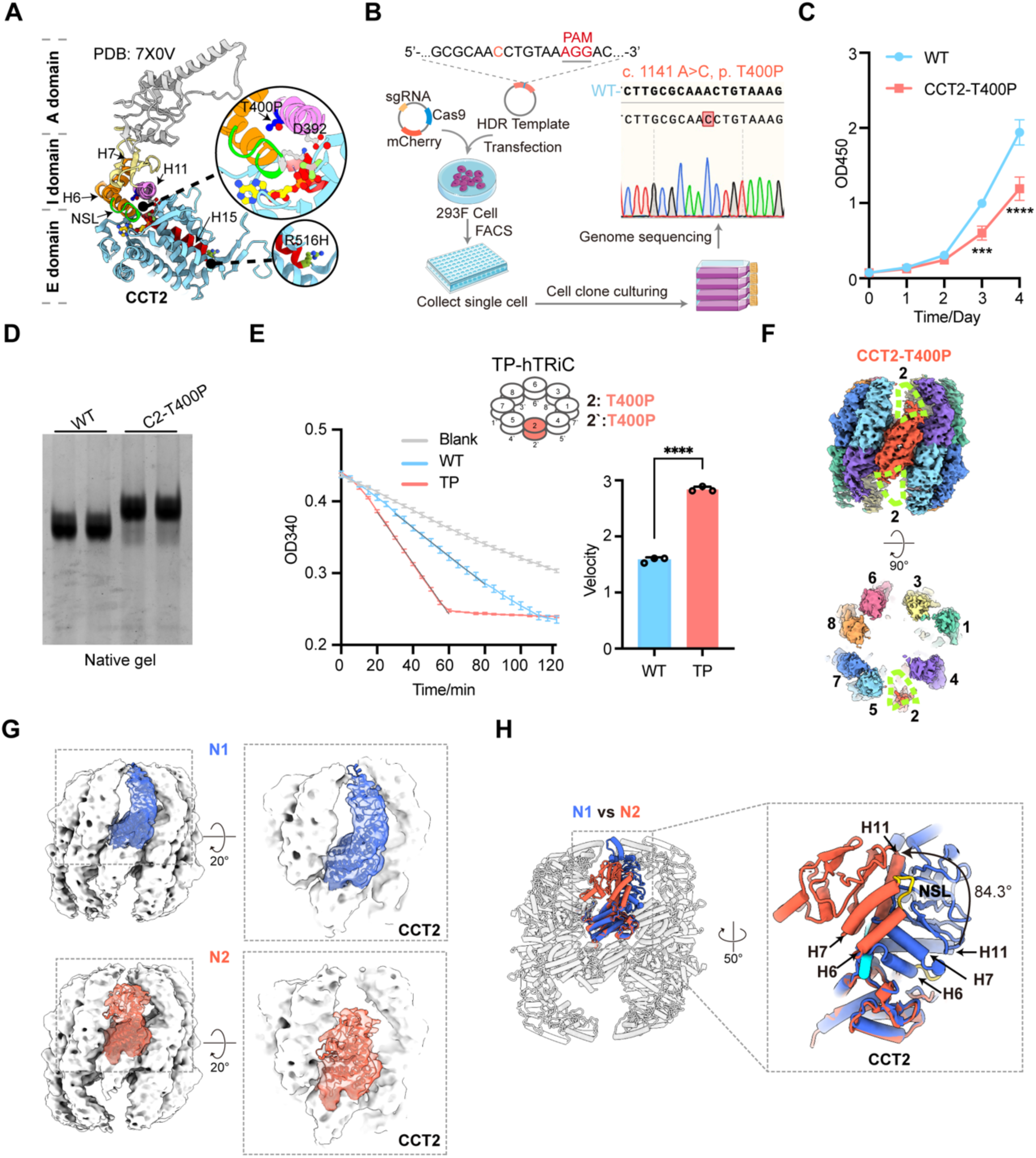
Construction of LCA-associated hTRiC mutants and analysis of altered structural dynamics. (A) Structural mapping of LCA-associated mutations on the human CCT2 subunit (PDB: 7X0V). Residue T400 is located on helix 11 (H11) in the I-domain, while R516 resides on helix (H15) at the C-terminus of the E-domain. The nucleotide-sensing loop (NSL) connecting helices H6 and H7 is also indicated. (B) Workflow for generating the CCT2-T400P mutant HEK293F cell line using CRISPR-Cas9 genome editing, accompanied by sequencing validation of the engineered cell line. (C) Cell viability of WT and CCT2-T400P 293F cells assessed by CCK-8 assay. Statistical analysis was performed using two-way ANOVA: ***p < 0.001, and ****p < 0.0001. (D) Coomassie blue-stained native-PAGE gel showing the slower migration of CCT2-T400P hTRiC compared to WT hTRiC, indicative of altered structural dynamics. (E) NADH-coupled ATPase assay, showing accelerated ATP-hydrolysis by the CCT2-T400P mutant hTRiC. ATPase rates are quantified in histogram. Statistical analysis was performed using one-way ANOVA: ****p < 0.0001. (F) Side and end-on views of the CCT2-T400P hTRiC consensus cryo-EM map in the NPP state. The missing density for the CCT2 A-/I-domain is outlined with a yellow-green dotted line. (G) Model-to-map fitting of the CCT2 subunit in the N1 (royal blue) and N2 (tomato) states. (H) Superposition of the N1- and N2-state CCT2 models indicates that the dominant movement in the N2 state originates from helices H6, H7, and H11 within the intermediate domain. The nucleotide sensing loop (NSL), which links helices H6 and H7, is highlighted in gold.

By integrating cryo-electron microscopy (cryo-EM), biochemistry, and proteomics, we elucidated the molecular basis underlying TRiC dysfunction in LCA. We engineered human HEK293F cell lines harboring the LCA-associated CCT2-T400P mutation and a yeast model carrying the CCT2-T394P/R510H double mutation (corresponding to human T400P/R516H). We show that these mutations disturb TRiC’s key intra-molecular and intra-ring allosteric networks, shifting the conformational equilibrium away from the folding-competent closed state. Beyond canonical substrates, our proteomic analysis identifies a potential role for TRiC in the biogenesis of SLC membrane transporters. We demonstrate that LCA-linked TRiC dysfunction leads to the downregulation of these transportors, which may be particularly catastrophic for the metabolically demanding tissues such as retina and embryo. These findings provide a structural basis for understanding how specific defects in the TRiC chaperonin lead to proteostasis imbalance and severe human disease.

## Results

### Construction of LCA-associated TRiC mutants and the altered biochemical properties

To elucidate the structural and functional impacts of LCA-associated mutations, we generated a homozygous CCT2-T400P HEK293F cell line using CRISPR-Cas9 gene-editing technology (Fig. 1B). The mutant cell line exhibited significantly reduced proliferation compared with wild-type (WT) cells (Fig. 1C), consistent with the embryonic lethality reported in homozygous T400P mice^18^. This growth defect suggests that the CCT2-T400P mutation severely compromises TRiC-mediated proteostasis essential for cell viability.

We next purified the CCT2-T400P mutant human TRiC (hTRiC) complex for functional characterization (Fig. S1C-D). Biochemical analysis revealed that the purified CCT2-T400P hTRiC migrated more slowly on native-PAGE than the WT complex, indicative of altered conformational dynamics (Fig. 1D). We further assessed the ATPase activity of WT and mutant hTRiC using a steady-state NADH-coupled ATPase assay. The CCT2-T400P mutant exhibited elevated ATPase activity relative to WT (Fig. 1E), indicating that the LCA-associated CCT2-T400P mutation significantly alter the ATP hydrolysis dynamics of TRiC. Since our attempt to generate a homozygous CCT2-R516H cell line were unsuccessful, we generated a complementary *S. cerevisiae* model harboring the homologous CCT2 mutations (T394P and R510H) to investigate the compound heterozygous effects in a eukaryotic system (Fig. S1E).

### The CCT2-T400P mutation induces profound structural flexibility in TRiC

To visualize the structural impact of the CCT2-T400P mutation, we performed cryo-EM analysis of the endogenously purified CCT2-T400P hTRiC complex in the absence of added nucleotide (Fig. S2A-C). We obtained a consensus map at 4.53 Å resolution, with the more stable E-domain at the local resolution of 4.0 Å (Fig. 1F, Fig. S2C-E). The overall architecture resembles the WT nucleotide partially preloaded (NPP) state, which contains endogenous ADP bound to the CCT3, CCT6, and CCT8 subunits^5,19^ (Fig. S2F-G). Notably, density corresponding to the A- and I-domains of CCT2 was strikingly absent in both rings, indicating that the T400P mutation induces substantial conformational dynamics in the CCT2 subunit (Fig. 1F).

3D variability analysis (3DVA) confirmed continuous, large-scale movement of the A- and I-domains of the CCT2 subunit pair (Fig. S2H). Further focused classification of the dynamic CCT2 subunit resolved three distinct conformational states (N1-N3, Fig. S2C). The N1 state resembles the WT NPP conformation (Fig. S2I). In contrast, the N2 state displays a disordered CCT2 A-domain and a pronounced outward rotation of the I-domain relative to the E-domain. Relative to the E-domain, the I-domain (containing helices H6, H7, and H11) undergoes an ∼84.3° rotation (Fig. 1G-H). Notably, this movement displaces the nucleotide-sensing loop (NSL) connecting helices H6 and H7 away from the E-domain (Fig. 1H, Movie. S1), suggesting a potential alteration in nucleotide-sensing capacity. In the N3 state, densities corresponding to the CCT2 A- and I-domains in the trans-ring remain absent (Fig. S2C, J). Together, these observations reveal that the CCT2-T400P mutation induces profound conformational dynamics within the TRiC complex, potentially modulating its allosteric regulation.

### CCT2-T400P disrupts the orchestration of TRiC conformational cycling

To examine how the T400P mutation perturbs the ATP-driven conformational cycle of TRiC, we determined cryo-EM structures of the CCT2-T400P mutant hTRiC in the presence of ATP-AlFx, an ATP-hydrolysis transition-state analog that stabilizes TRiC in the closed state^20,21^ (Fig. S3A-C). We resolved four distinct conformational states (S1-S4; Fig. 2A-D): (i) a predominant open state at 3.03 Å resolution (S1), with ATP bound in all subunits and an attacking water molecule observed in CCT2, CCT6, and CCT8 (Fig. S4I); (ii) a previously uncharacterized intermediate state at 3.43 Å resolution (S2), characterized by increased separation between the two on-axis CCT2 subunits at the E-domain, resembling a slightly less open comformation; and (iii) two double-ring closed states: an empty TRiC complex at 3.28 Å resolution (S3) and a substrate-containing complex at 3.23 Å resolution (S4) (Fig. S3D, Fig. S4A-G).

**Fig. 2.**
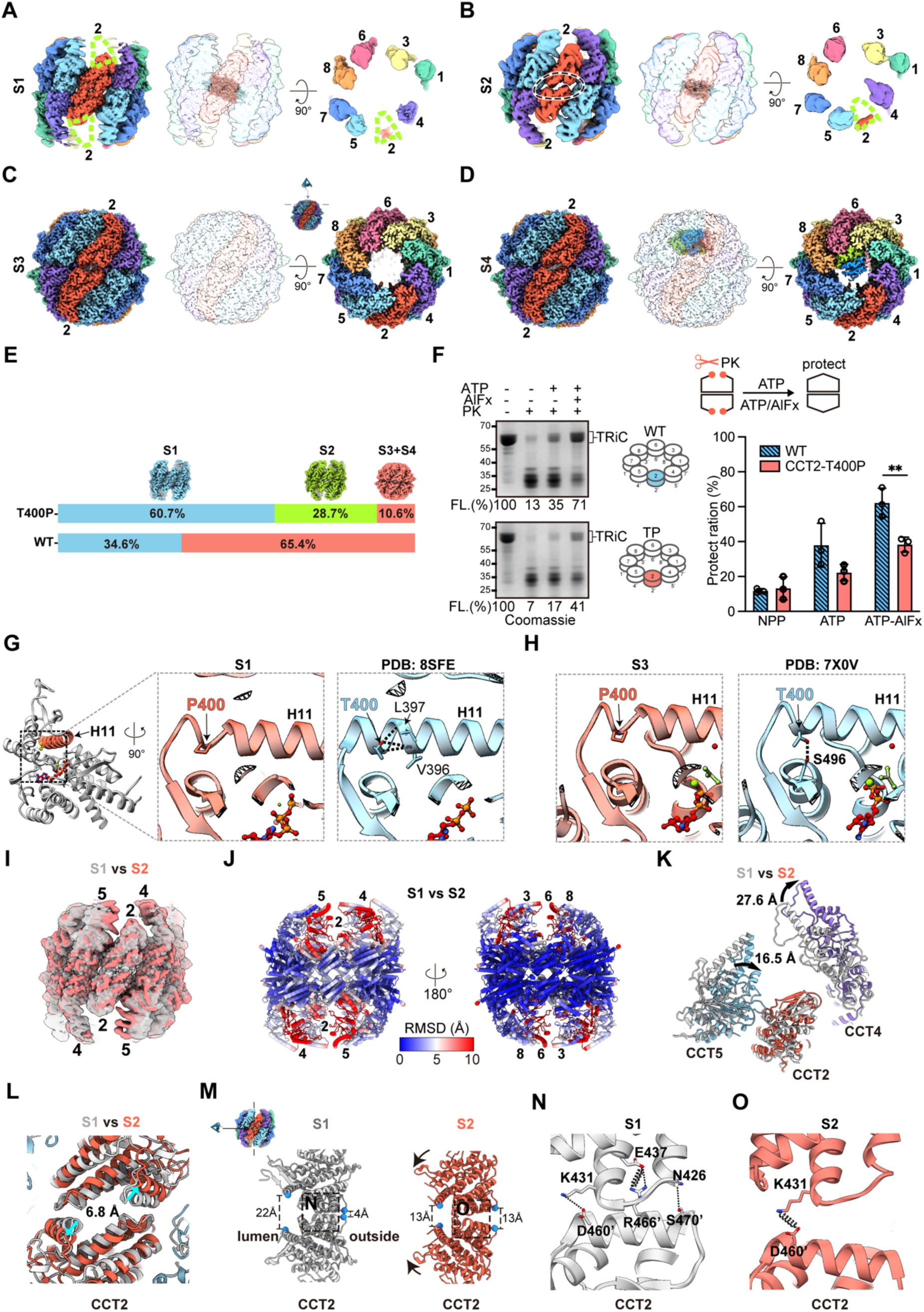
CCT2-T400P disrupts the orchestration of hTRiC conformational cycling. (A) Side and end-on views of the CCT2-T400P hTRiC in the ATP-AlFx open S1 state. The missing density for the A-/I-domains is outlined with a yellow-green dotted line. (B) Cryo-EM map of the intermediate S2 state; the missing density for the A-/I-domain is highlighted in the end-on view. (C-D) Cryo-EM maps of CCT2-T400P TRiC in the closed, ATP-hydrolysis transition state with an empty (C, S3) or substrate-filled (D, S4) chamber. (E) Population distribution comparasion between the mutant hTRiC and the available WT hTRiC both in the presence of ATP-AlFx. (F) Coomassie blue-stained SDS-PAGE and statistical analysis of the proteinase K (PK) protection assay. ATP induces conformational cycle, while ATP-AlFx stablizes the PK-protected closed state. The percentage of full-length TRiC remaining after PK digestion is annotated below each lane and quantified in histograms (n=3). Statistical analysis was performed using one-way ANOVA: **p < 0.01. Reduced PK protection is observed for the CCT2-T400P variants. (G) Comparison of H-bond networks at the site 400 within H11 in the mutant versus WT CCT2 (PDB: 8SFE) in the open state. (H) In WT CCT2 (PDB: 7X0V, right), a H-bond between T400 and S496 exists, which may contribute stabilizing the closed conformation; while the T400P mutation disrupts this interaction. (I) Overlay of CCT2-T400P mutant maps in the S1 (gray) and S2 (Tomato) states, viewed from the CCT2 side. (J) RMSD analysis between the S1 and S2 states highlights significant movement of subunits on CCT2 side, while subunits on the CCT6 side remain stationary. (K) Rotation distance of CCT4 and CCT5 during the transition from S1 to S2. (L) Visualization of the CCT2 pair separating from the E domain. (M) Structural models of the CCT2 pair in the S1 and S2 states. Distances are measured between the E471-E471’ and E22-E22’ residues. Detailed inetarction networks within the black dotted squares are expanded in (N) and (O). (N) H-bond (dotted line) and salt bridge (spring) interactions within the CCT2 pair in the S1 state. (O) Salt bridge interactions within the CCT2 pair in the S2 state.

Particle distribution analysis revealed a profound shift in the conformational landscape. The mutant complex predominantly populated the open S1 state (60.7%) and the less-open intermediate S2 state (28.7%), with only a minor fraction reaching the closed S3 and S4 states (10.6%; Fig. 2E). This distribution contrasts sharply with that of WT hTRiC, which is predominantly closed (65.4%) under the same conditions (Fig. 2E)^19^. We further validated these structural observations using a proteinase K (PK) protection assay to assess TRiC’s ability to undergo ring closure during ATP hydrolysis. This assay distinguishes between the PK-sensitive open state and the PK-resistant closed state of TRiC^20,22^. Whereas WT hTRiC exhibited robust PK resistance in the presence of ATP-AlFx (>60% intact), the CCT2-T400P mutant showed significantly reduced PK protection (∼40% intact) (Fig. 2F), corroborating the significally reduced closed-state population observed by cryo-EM (Fig. 2E). Collectively, these structural and functional data demonstrate that the T400P mutation compromises TRiC’s ability to adopt the functionally critical closed state, likely by disrupting allosteric coordination among subunits.

### T400P mutation alters CCT2 A-/I-domain dynamics and key atomic interactions

Structural comparison of the mutant and WT TRiC provides a molecular explanation for the reduced closed-state population. In the ATP-bound open S1 state, the overall architecture of the mutant complex closely resembles that of WT TRiC (Fig. S5A), except for the CCT2 A-/I-domains, which remain highly dynamic and unresolved (Fig. 2A). This contrasts with human and bovine WT TRiC, in which ATP binding stabilizes the otherwise dynamic CCT2 subunits^7,19^ (Fig. S3E). This suggests that the T400P mutation potentially disrupts the allosteric network between CCT2 A-/I-domains and the ATP-binding E-domain, thereby impairing coordinated conformational changes upon nucleotide binding. Locally, the T400P substitution in helix H11 perturbs main-chain hydrogen-bond networks, likely increasing the flexibility of H11 (Fig. 2G, Fig. S5C), which caps the nucleotide-binding pocket.

Although the closed, empty S3 state resembles the WT closed conformation (Fig. 2C, Fig. S5B), the T400P mutation abolishes a key intramolecular H-bond between T400 and S496, located adjacent to the nucleotide-binding pocket (Fig. 2H, Fig. S5D). This may potentially destabilize the closed conformation and promoting a premature return to the open state (Fig. 2E).

Furthermore, the unique intermediate S2 state captures a snapshot of the TRiC ring closure process (Fig. 2B, I, Fig. S5E). In this state, the CCT2-side of the ring (CCT5/CCT2/CCT4) pivots inward (Fig. 2J), with CCT5 moving toward CCT2 by ∼16.5 Å, CCT4 moving toward CCT1 by ∼27.6 Å (Fig. 2K), and CCT2 shifting ∼6.8 Å away from the E-domain (Fig. 2L). Meanwhile, the opposite side of the ring (CCT3/CCT6/CCT8) remains in an open-like conformation (Fig. 2J), reminiscent of the asymmetric transition state observed in the TRiC-Gβ-PHLP1 complex^23^. This stepwise movement is coordinated by remodeling of the inter-ring interface at the on-axis CCT2-CCT2’ subunit paris. Relative to the open state, the spacing between the outer E471-E471’ residues increases from 4 Å to 13 Å, whereas the distance between the inner E22-E22’ residues decreases from 22 Å to 13 Å (Fig. 2M). At this interface, the conserved inner K431-D460’ salt bridge acts as a pivot point (Fig. 2N-O, Fig. S5F-H). Although obtained from mutant complex, this stepwise, asymmetric mechanism may provide insight into the native TRiC ring-closure process.

### TRiC facilitates α-tubulin folding via a unique interaction with the CCT8 C-terminal tail

Symmetry expansion and focused classification of the substrate-bound closed S4 TRiC yielded a 3.45-Å-resolution map, revealing a co-purified, near-native α-tubulin within TRiC chamber (Fig. 3A-C, Fig. S3C, S4G-H). This identity was comfirmed by mass spectrometry (Table. S2) and by distinctive structural features, particularly the distinct S9-10 loop, which is eight residues longer (aa 364-371) in α-tubulin than in β-tubulin (Fig. 3D, Fig. S6). This structure represents the first direct visualization of TRiC folding α-tubulin, extending the established structural framework for TRiC-mediated β-tubulin folding^19,24-26^. Mass spectrometry showed slightly higher abundance of the α-tubulin isoform TUBA1C compared with TUBA4A (Table. S2). Given the high sequence identity between these isoforms (Fig. S6), TUBA1C was used for structural modeling. The α-tubulin appears fully folded, as evidenced by well-resolved density for bound GTP (Fig. 3B-C) and for the typically flexible K40 loop (aa 37-46) (Fig. 3E), which is unsolved in prior microtubule structures^27-29^. These feature undersore the role of TRiC in stabilizing the final native fold of α-tubulin. Importantly, the CCT2-T400P mutant retains this folding capability, positioning α-tubulin within the TRiC chamber in a position similarly to β-tubulin (Fig. S7A-B) via interactions with the CCT1/3/6/8 hemisphere via hydrogen bonds and salt bridges (Fig. S7D-F, Table. S3).

**Fig. 3.**
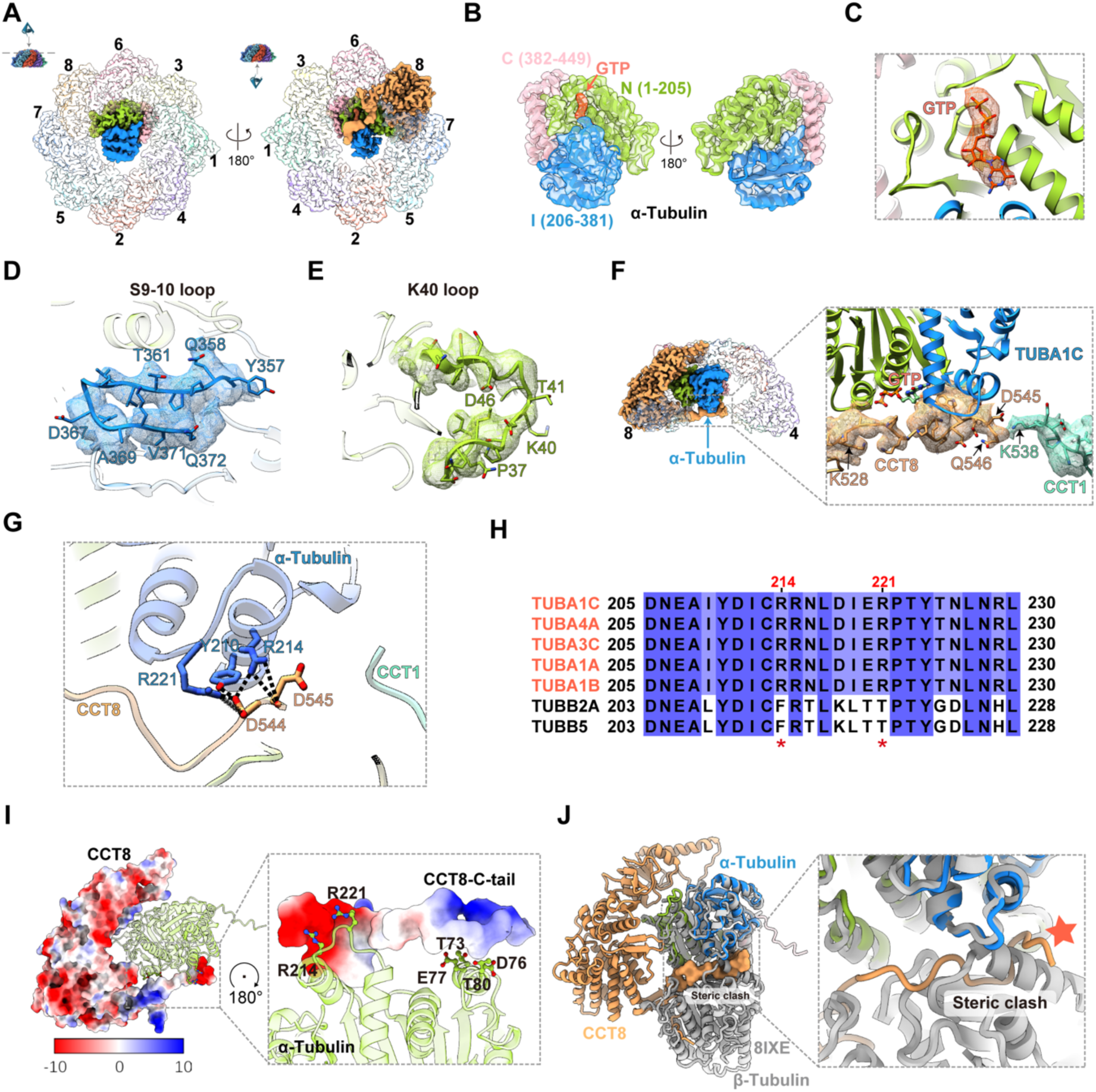
Near-native state α-tubulin in the TRiC chamber. (A) Cryo-EM map of the CCT2-T400P TRiC in the closed state with encapsulated α-tubulin (top and bottom views). The CCT8 subunit is highlighted in sandy brown, showing its C-terminal tail cradling α-tubulin within a single ring. (B) Map-model fitting of the nearly fully folded α-tubulin (TUBA1C) inside the TRiC chamber, featuring bound GTP. The N-domain is colored yellow-green, the I-domain dodger blue, and the C-domain light pink. (C) Detail of the bound GTP in α-tubulin. (D) Resolved density for the α-tubulin S9-10 loop. (E) Density for the stabilized K40 loop of α-tubulin within the TRiC chamber. (F) The CCT8 C-terminal tail inetracts with α-tubulin near its GTP-binding pocket and contacts the C-terminus of CCT1; this CCT8-CCT1 connection isolates α-tubulin within the ring. (G) H-bond and salt bridge interactions between the CCT8 C-tail and α-tubulin. (H) Sequence alignment of α- and β-tubulin showing that key residues (R214, R221) mediating the interaction with the CCT8 C-tail are conserved in α-tubulin but not in β-tubulin. (I) Electrostatic surface potential analysis demonstrates charge complementarity between the CCT8 C-tail and its binding site on α-tubulin. (J) Superimposition of the CCT8-α-tubulin conformation with the α/β-tubulin dimer (PDB: 8IXE) reveals a steric clash between the CCT8 C-tail and β-tubulin, suggesting a mechanism for preventing premature dimerization.

Strikingly, α-tubulin folding involves a unique interaction with CCT8 that is not observed in previous TRiC-β-tubulin structures. The flexible CCT8 C-terminal tail extends to cradle α-tubulin near its GTP-binding pocket, stretching to contact CCT1 C-terminus to anchor α-tubulin (Fig. 3F). This interaction appears specific for α-tubulin (Fig. S7C)^19^. The α-tubulin residues Y210, R214, and R221 form hydrogen bonds and salt bridges with the CCT8 C-terminal tail (Fig. 3G), further stabilized by electrostatic contacts between α-tubulin helix 73-80 and the CCT8 C-terminal tail (Fig. 3I). Notably, the highly conserved R214 and R221 are unique to α-tubulin (Fig. 3H), providing the molecular basis for CCT8’s selective folding. The CCT8 N-terminal tail from the opposite ring also contact α-tubulin (Fig. S7G), consistent with interactions observed in β-tubulin^19^. Moreover, the CCT8 C-terminal tail occupies the αβ-tubulin dimerization interface, potentially preventing premature αβ-tubulin association in TRiC chamber (Fig. 3J). Together, these findings highlight the versatility of TRiC’s unstructured tails in orchestrating substrate-specific folding pathways.

### Disrupted TRiC intra-ring coordination revealed by yeast CCT2-T394P-R510H mutant

To investigate the compound heterozygous effects of LCA-associated mutations, we constructed and analyzed a yeast TRiC (yTRiC) harboring the analogous CCT2 mutations T394P and R510H (Fig. S1F-G). Cryo-EM analysis of the NPP dataset yieled two yTRiC conformational states (yTRiC-NPP-Y1, -Y2; Fig. S8A-F), both displaying the characteristic Z-shaped CCT2 subunit pair and ADP bound to CCT3/CCT6/CCT8 (Fig. 4A, Fig. S8G-H). Y1 resembles WT yTRiC (PDB: 5GW4), and Y2 exhibits an inward-tilted CCT7 (Fig. 4A). A similar Y2-like conformaiton can also be detected upon reanalysis of the original WT yTRiC NPP state dataset^5^ (data not shown). More importantly, and consistent with observations in hTRiC, in the presence of ATP-AlFx, cryo-EM analysis revealed a significantly reduced closed-state population in the double mutant (53.4% closed) compared with WT yTRiC, which is almost all closed under the same condition^21,30^ (Fig. 4B-C, Fig. S9A-C). Despite this defect, the overall architecture of the mutant complex closely resembles that of the WT (Fig. 4D, Fig. S9H-I).

**Fig. 4.**
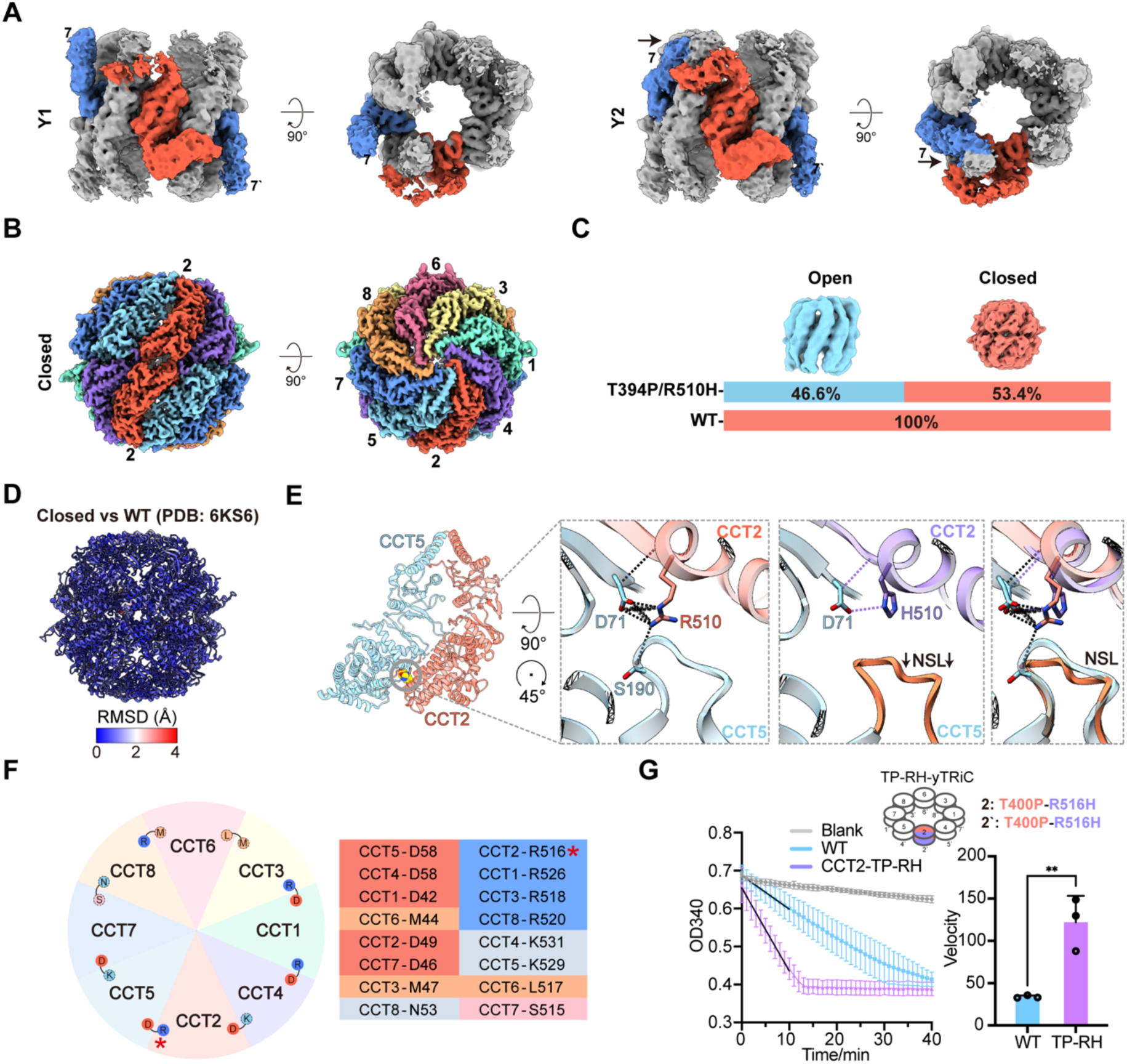
Structural analysis of the CCT2-T394P-R510H yTRiC. (A) Cryo-EM maps of the CCT2-T394P-R510H yTRiC in the NPP state: Y1 state showing an outward-tilted CCT7, and Y2 state showing an inward-tilted CCT7. (B) Side and end-on views of the closed-state map of the CCT2-T394P-R510H yTRiC reconstructed from the ATP-AlFx dataset. (C) Population distributions of the open and closed states in the mutant versus the WT dataset. (D) RMSD analysis comparing the closed state of the CCT2-T394P-R510H yTRiC to the WT yTRiC (PDB: 6KS6). (E) Detailed analysis of the interface between CCT2 residue 510 and CCT5. The CCT2-R510H mutation disrupts H-bond and salt bridge interactions, causing the CCT5 NSL to move away from CCT2. (F) Diagram illustrating the conserved R-D interaction between adjacent TRiC subunits, which are essential for ring opening and closure in Group II chaperone. These R-D interaction pair are conserved in CCT2-CCT5, CCT1-CCT4, and CCT3-CCT1 interfaces, but vary in other subunits as depicted. (G) NADH-coupled ATPase assay showing accelerated ATP hydrolysis by the CCT2-T394P-R510H yTRiC mutant. ATPase rates are quantified in the histograms. Statistical analysis was performed using one-way ANOVA: **p < 0.01.

Detailed structureal analysis implicates the R510H mutation in disrupting intra-ring allostery. In WT yTRiC, CCT2-R510 forms a critical H-bond with CCT5-S190 in the nucleotide-sensing loop (NSL), as well as a salt bridge and H-bond network with CCT5-D71 (hereafter referred to as the R-D interaction; Fig. 4E, Fig. S9D). This R-D interaction is conserved across group II chaperonins in both open and closed states^31^ (Fig. S9G), and across multiple subunit interfaces (e.g., CCT2-CCT5, CCT1-CCT4, and CCT3-CCT1), where it functions as a linchpin for coordinated ring movements (Fig. 4F, S9J-M). The R510H mutation abolishes the H-bonds with CCT5-S190, causing the CCT5 NSL to shift away from CCT2 (Fig. 4E), potentially impairing coordinated CCT2-CCT5 movement in TRiC’s allosteric cycle. Together, the R510H mutation weakens these key stabilizing contacts, uncoupling the coordinated movement between CCT2 and CCT5 and impairing efficient ring closure. Consistent with these structural defects, the LCA-associated mutant yTRiC exhibits elevated ATPase activity (Fig. 4G), mirroring the deregulation observed in the in the human CCT2-T400P mutant (Fig. 1E), suggesting a significantly altered ATP hydrolysis dynamics in TRiC.

### CCT2-T400P mutation alters the proteome and impairs membrane transporter biogenesis

To elucidate the cellular consequences of TRiC dysfunction, we performed whole-cell label-free quantitative mass spectrometry to compare the proteomic profiles of WT and CCT2-T400P HEK293F cells. We identified 180 significantly downregulated and 57 upregulated proteins in the mutant cells (Fig. 5A). Notable upregulated proteins include CAPG, an actin-capping gelsolin linked to retinal injury responses^32^, and ALDH1A1, a key retinoic acid signaling enzyme^33^, indicating a compensatory response to metabolic stress. Conversely, OXCT1, a critical mitochondrial ketone body catabolic enzyme^34^, was the most downregulated protein, indicative of impaired cellular metabolism. Additional downregulated proteins included several solute carrier (SLC) transporter family proteins (SLC16A7, SLC7A11, and SLC38A2), indicating membrane transport disruption, and ubiquitin pathway components (FBXW9, UBE2C, and WDTC1), pointing to altered protein degradation and overall proteostasis. Notably, TRiC subunits themsevles was unchanged or only slightly downregulated, indicating that the CCT2-T400P mutation does not prevent the complex from assembly but likely impairs its function.

**Fig. 5.**
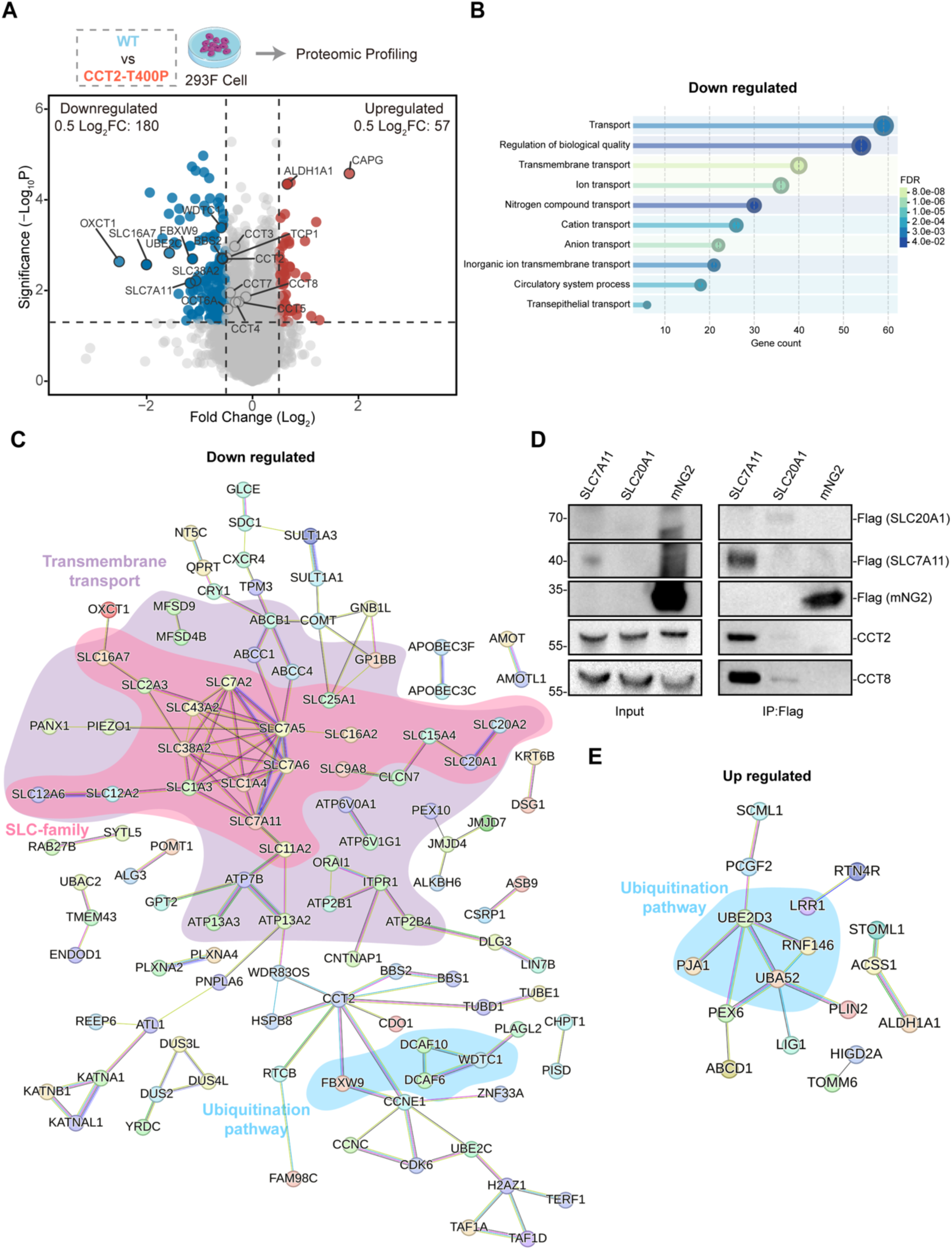
LCA-associted mutations alter the cell proteomic profiles. (A) Workflow for label-free quantificative proteomics and volcano plot showing differentially abundant proteins in CCT2-T400P versus WT HEK293F cells. Dashed lines indicate thresholds for log2 fold change (≥0.5) and significance (P ≤ 0.05). Significantly upregulated proteins are shown in red, and downregulated proteins in blue. TRiC subunits and key proteins discussed in the manuscript are labeled. (B) GO enrichment analysis of downregulated proteins reveals predominance of biological processes related to transport. (C) PPI network of downregulated proteins highlights clusters associated with transmembrane transport (purple), particularly SLC family members (Pink). Several WD40 repeat-containing proteins previously reported to interact with TRiC and function in the ubiquitin pathway (blue) were also identified. (D) Co-immunoprecipitation of FLAG-tagged transporters SLC7A11 and SLC20A1 confirms their interaction with endogenous TRiC subunits CCT2 and CCT8. FLAG-tagged mNG2 served as a negative control (n=2). (E) Upregulated proteins shows no distinct GO term enrichment; several upregulated proteins associated with the ubiquitin pathway are highlighted in blue.

Gene Ontology (GO) analysis of downregulated proteins revealed a significant impact on transmembrane transport (Fig. 5B). The SLC superfamily, critical for ions and metabolite transport, often coupled with ion gradients such as Na⁺ or H⁺ ^35^, was broadly reduced (Fig. 5C). This included SLC7A11 (also known as xCT), which is essential for neuroprotection against oxidative stress and cell proliferation during development, as well as SLC38A2, a sodium-coupled neutral amino-acid transporter^36,37^. Both of these transporters are likely involved in regulating retinal metabolism. Co-immunoprecipitation confirmed that two members of SLC transporters, SLC7A11 and SLC20A1 interact with TRiC (Fig. 5D), suggesting that they may be potential TRiC clients. Similarly, several ABC transporters, which are implicated in inherited retinal diseases, were downregulated (Fig. 5C). Components of the ubiquitination pathway were also dysregulated (Fig. 5C, E). Together, these data uncover a previously unrecognized role for TRiC in the biogenesis of membrane transporters, and suggest that impaired proteostasis of these transporters may contribute to retinal degeneration in LCA and potentially to embryonic developmental defects (discussed later).

## Discussion

Although CCT2 mutations T400P and R516H are known drivers of LCA, their underlying pathogenic mechanisms remain elusive. Using complementary human and yeast models, we show that these mutations disrupt TRiC’s allosteric network and its productive conformational cycle. Specifically, the mutants rarely populate the folding-active closed state, with closed population reduced from 65.4% to 10.6% in human system and from 100% to 53.4% in yeast (Fig. 2E, 4C). This structural instability, evidenced by increased protease susceptibility (Fig. 2F), is coupled with accelerated yet futile ATPase activity (Fig. 1E, 4G). This indicats a wasteful, energy-consuming cycle that fails its biological mission. This functional collapse significantly impairs cellular proliferation in the homozygous CCT2-T400P cells (Fig. 1C), providing a mechanistic explanation for the embryonic lethality observed in CCT2-T400P mouse models^18^.

The chaperonin cycle relies a fine-tuned equilibrium between substrate-accepting open states and folding-active closed states^38^. Our data demonstrate that LCA-associated mutations drastically shift this balance toward open states (Fig. 2E, 4C), precluding the formation of the closed chamber required for folding. Our high-resolution cryo-EM structures pinpoint the atomic determinants of this dysfunction. The CCT2-T400P mutantion disrupts a critical intramolecular H-bond (T400-S496) at the I-E domain interface, destabilizing the closed conformation (Fig. 2H). Complementarily, the yeast model shows that the CCT2-R510H mutation (analogous to human R516H) disturbs a conserved intensive salt bridge/H-bounds network between CCT2 and CCT5 (Fig. 4E), essential for intra-ring allosteric coordination required for ring closure. These results demonstrate how two distinct pathogenic mutations converge to disrupt the chaperonin cycle by distorting conserved intra-molecular and intra-ring interactions. Such findings highlight the exquisite evolutionary optimization of TRiC and offer a structural explanation for its role in LCA pathogenesis.

Unexpectedly, our structural data captured a folded α-tubulin within the mutant hTRiC chamber, uncovering a folding mechanism distinct from that of β-tubulin (Fig. 3). Here, the flexible C-terminal tail of CCT8 forms a unique cradle that stabilizes α-tubulin intermediates and likely provides steric shielding against premature αβ-tubulin dimerization. These results highlight a broader principle: TRiC’s disordered C-terminal tails functions as specialized affinity clamps that contribute to the biogenesis of key cytoskeletal components.

Our proteomic analysis further revealed an unexpected requirement for TRiC in the biogenesis of membrane transporters (Fig. 5C). The selective downregulation of SLC and ABC transportors in CCT2-T400P cells, coupled with evidence of their interactions with TRiC (Fig. 5D), suggests an unpresedent pathological axis. We propose that impaired folding of these essential transporters may represent the primary driver of the developmental disorders and embryonic lethality linked to the T400P mutation^18^.

These substrate-specific defects provide a compelling explanation for the genotype-phenotype correlations observed in murine models. While CCT2-T400P and R516H mutations were initially identified as compound heterozygotes in LCA patients, subsequent murine models revealed a starking divergence in phenotypic severity: T400P is embryonically lethal, while R516H results in a retina-specific phenotype^18^. This suggests that the R516H allele retains sufficient chaperonin activity to sustain embryogenesis, effectively rescuing the T400P-associated lethality in compound heterozygous patients. While retinal degeneration in R516H mice has been attributed to the depletion of ciliary proteins, particularly CCDC181, our data indicate that the more severe T400P mutation causes a broader collapse of proteostasis that extends to essential membrane transporters. Consequently, we propose that ciliary dysfunction drives the retina-specific LCA phenotype, while widespread transporter disruption underlies the systemic developmental defects and embryonic lethality.

Previous reports proposed that LCA-related CCT2 mutations may stabilize the TRiC closed state by reducing the ADP-off rate, hindering the release of monomeric CCT2 for non-canonical autophagic functions^39,40^. However, our structural and biochemical evidence consistently points to the opposite: a destabilized closed state and a disrupted holo-complex cycle. Although these discrepancies may arise from differing experimental contexts, our data suggest that LCA pathology primarily stems from the loss of TRiC holo-complex chaperonin activity. Whether dysfunction of monomeric CCT2 also contributes to the disease phenotype remains a subject for further study.

In summary, we have identified the atomic basis for how LCA-associated mutations disrupt TRiC allosteric network and functional cycling by destabilizing essential intra-molecular and intra-ring contacts. This work provides a mechanistic links between chaperonin mechanics and a specific class of cellular substrates, membrane transportors, thereby establishing a new framework for understanding the etiology of TRiCopathies and congenital retinal disorders.

## Methods and materials

### Construction of the CCT2-T400P HEK293F cell line

We generated a HEK293F cell line harboring a homogeneous CCT2-T400P mutation using CRISPR/Cas9 genome editing system. A single-guide RNA (sgRNA, TGTTCTTGCGCAAACTGTAA) was designed and cloned into the pX330-Cas9 expression vector, which included a mCherry fluorescent marker for tracking transfection efficiency and enabling fluorescence-activated cell sorting (FACS). A donor template containing the CCT2-T400P mutation and homology arms was amplified from the HEK293F genome via PCR and cloned into a pGEX-T vector for homology-directed repair (HDR). The donor template and the pX330-sgRNA-mCherry vector were co-transfected into HEK293F cells using polyethyleneimine as the transfection reagent. After 48 h, mCherry-positive cells were isolated into 96-well plate using FACS, and cultured for 14-21 days to allow clonal expansion. Genomic DNA was extracted (Mouse Direct PCR Kit, Selleck, B40013), and the targeted region was amplified and validated by Sanger sequencing to confirm the homogeneous CCT2-T400P mutation.

### Construction of CCT2-T394P-R510H yeast strain

The *S. cerevisiae* CCT2-T394P-R510H mutant was constructed using a two-step cloning strategy. First, the *CCT2* gene and its downstream region (*CCT2* Down) were PCR-amplified from yeast genome, and coloned into linearized pUC19 vectors with a HIS3 selection marker. The T394P and R510H mutations were introduced by site-directed mutagenesis (Fig. S1E). Second, the linearized fragment (CCT2-T394P-R510H-HIS3-downstream) was amplified and transformed into the haploid yeast strain carrying a Strep-CBP-His-tagged CCT3 subunit^5^ for affinity purification. Transformants were selected on histidine-deficient media, and positive clones were confirmed by sequencing.

### Purification of human TRiC

Human TRiC (hTRiC) was purified as previously described^19^ with minor modifications. Briefly, WT or CCT2-T400P HEK293F cell pellets were lysed in MQA buffer (20 mM HEPES pH 7.4, 50 mM NaCl, 5 mM MgCl_2_, 0.1 mM EDTA, 1 mM DTT, 10% glycerol, 1 mM PMSF, and 1 mM ATP) with protease inhibitor (Roche). Cell lysates were clarified (18,000 rpm, 30 min) and ultracentrifuged (150,000 ×g, 1.5 h). The cytoplasmic fraction was filtered through a 0.45 μm membrane and loaded onto a Q Sepharose column (Cytiva). TRiC was eluted with a 10–80% gradient of MQB buffer (MQA with 1 M NaCl) containing 5% glycerol. TRiC-containing fractions were pooled and diluted with MQA to ∼100 mM NaCl, and purified via HiTrap Heparin HP chromatography (Cytiva). TRiC was again eluted using a 10–80% MQB gradient. TRiC-containing fractions were pooled and incubated with 10 mM ATP at 37 °C for 30 min to promote substrate release, then concentrated and subjected to size-exclusion chromatography (SEC) on a Superose 6 Increase 10/300 GL column (Cytiva) in ATP-free MQA buffer containing 5% glycerol. TRiC was eluted in the fraction range of 12.5–15 mL, which is consistent with the expected size of a ∼1 MDa complex. Finally, multiple rounds of buffer exchange were performed to remove residual ATP, yielding biologically active TRiC.

### Purification of yeast TRiC

Yeast TRiC (yTRiC) was purified as previously described^5^. Briefly, the supernatant from WT or mutant yeast lysate was incubated with Calmodulin Sepharose (Cytiva) overnight at 4°C. The complex was eluted in buffer containing 20 mM Hepes pH 7.5, 5 mM MgCl_2_, 0.1 mM EDTA, 50 mM NaCl, 1 mM DTT, 10% glycerol, and 2 mM EGTA. The pooled eluates containing TRiC were further purified by SEC (Superose 6 Increase, Cytiva). Peak fractions containing TRiC were concentrated using a Millipore Ultrafree centrifugal filter device (100 kDa cutoff).

### Cryo sample preparation and data acquisation

For nucleotide-partially-preloaded (NPP) TRiC, purified TRiC was applied to a poly-Lysine pretreated holey carbon grid (Quantifoil, R1.2/1.3, 200 mesh), and vitrified using a Vitrobot Mark IV (Thermo Fisher Scientific). For the TRiC-ATP-AlFx sample, TRiC was incubated with 5 mM MgCl_2_, 5 mM Al (NO_3_)_3_, 30 mM NaF, and 1 mM ATP at 37 °C (hTRiC) or 30°C (yTRiC) for 1 h prior to freezing.

Images were acquired on a Titan Krios transmission electron microscope (Thermo Fisher Scientific) operated at 300 kV. The mutant yTRiC-NPP dataset was collected on a K2 Summit direct electron detector (Gatan) in super-resolution mode, with a pixel size of 1.318 Å/pixel. The exposure time for each frame was 0.2 s, and the total accumulation time was 7.6 s, with a total accumulated dose of 38.0 e^−^/Å^2^ on the specimen. Images were collected by utilizing SerialEM^41^. All other datasets were recorded on a K3 detector (Gatan) in counting mode utilizing EPU software (Thermo Fisher Scientific). For mutant-hTRiC-NPP and mutant-yTRiC-ATP-AlFx samples, the pixal size is 1.093 Å/pixel, and the accumulated dose is 50.2 e^−^/Å^2^ on the specimen. For mutant-hTRiC-ATP-AlFx sample, the pixel size is 0.86 Å, and the accumulated dose is 56.8 e^−^/Å^2^.

### Image processing and reconstruction

Single-particle analysis was performed using RELION 3.1^42^ and cryoSPARC 4.2.1^43^. All images were aligned and summed using MotionCorr2^44^. After CTF estimation using CTFFIND4^45^, particle auto-picking was performed utilizing crYOLO 1.7.6^46^.

For the CCT2-T400P-hTRiC-NPP dataset, 2D classification followed by two rounds of heterogeneous refinement using EMDB-32922 as the initial model yieled 107,501 particles with well-defined features. After subsequent CTF refinement and Bayesian polishing, we obtained a consensus map at the resolution of 4.53 Å. A subsequent 3DVA analysis^47^ revealed continuous movement in the CCT2 subunit pair (Fig. S2H). We then performed focused 3D classification on CCT2 using a broader mask, followed by non-uniform refinement, resolving three conformational states (N1, 4.81 Å; N2, 5.58 Å; and N3, 5.86 Å, Fig. S2C-E).

For the CCT2-T400P-hTRiC-ATP-AlFx dataset, 1,907,580 particles were initially picked from 10,842 micrographs. Following 2D classification, 3D classification in RELION, and heterogeneous refinement in cryoSPARC, using EMDB-32922 (open) and EMDB-32926 (closed) as initial models, 1,261,932 particles remained with open feature, and 235,789 particles with closed feature. The 1,261,932 particles was further separated by 3DVA-guided heterogeneous refinement, followed by CTF refinement, Bayesian polishing, and non-uniform refinement, yielding maps at 3.03 Å (S1, 598,570 particles) and 3.43 Å (S2, 282,675 particles). The closed fraction was subjected to focused classification based on internal chamber occupancy. Empty closed particles were reconstructed to 3.28 Å (S3, 62,780 particles). Substrate-bound particles were refined to 3.23 Å (S4, 41,331 particles). To improve the substrate density within the chamber, particles were refined with C2 symmetry, symmetry-expanded, and subjected to masked 3D classification focused on the half-ring to isolate optimal substrate features. A final subset of 25,657 particles with the clearest substrate density was reconstructed to a 3.45 Å resolution map. All final maps are sharpened by EMready^48^ (Fig. S3C-D).

For the CCT2-T394P-R510H-yTRiC-NPP dataset, 220,092 particles were picked from 1,210 micrographs. Following 2D and 3D classification, 94,016 particles were selected for heterogeneous refinement in cryoSPARC (using EMDB-9540 as the initial model), yielding 74,038 high-quality particles that produced a consensus map at 3.71 Å resolution. 3DVA analysis revealed continuous movement in the CCT2 and CCT7 subunits. Subsequent 3DVA clustering separated the particles into four groups, representing two distinct states of the complex. Non-uniform refinement of these subsets yielded reconstructions of the two states at 3.84 Å (41,850 particles) and 3.80 Å (32,188 particles) (Fig. S8A-F).

For the CCT2-T394P-R510H-yTRiC-ATP-AlFx dataset, 201,104 particles were initially picked. After 2D classification, 157,439 high-quality particles were selected for 3D classification (EMDB-0758 as the initial model). Subsequent heterogeneous refinement in cryoSPARC separated the particles into open and closed states (Fig. S9C). Following two additional rounds of heterogeneous refinement, 67,546 particles were selected to generate a final closed-state map at 3.92 Å resolution. A distinct subset of 58,871 particles exhibiting an open conformation was reconstructed to 7.32 Å resolution (Fig. S9E-F).

### Model building

Atomic models for the mutant human and yeast TRiC complexes were built using available TRiC structures (human: PDB 7X0A, 7X3J, 7X0S, 7X0V; yeast: PDB 5GW4, 6KS6, 6KS7) as templates. The human α-tubulin TUBA1C model was generated using AlphaFold3^49^. Point mutantations were introduced into the CCT2 subunit using COOT^50^. Individual subunits were docked into the cryo-EM density maps via rigid-body fitting in UCSF Chimera^51^, followed by flexible fitting with Rosetta^52^. Models underwent iterative refinement involved manual adjustment in COOT and phenix.real_space_refine in Phenix^53^. Final models were validated using Phenix. Figures and electrostatic surface property calculations were generated using UCSF Chimera and ChimeraX^54^. Interface analyses were performed using the PISA server^55^.

### Proteinase K protection assay

Purified TRiC complexes were diluted to a final concentration of 0.3 μM in MQA buffer (20 mM HEPES pH 7.5, 50 mM NaCl, 5 mM MgCl_2_, 0.1 mM EDTA, 1 mM DTT, 5% glycerol). Digestion was performed with Proteinase K (PK; Sigma-Aldrich, P2308) at a final PK concentration of 20 μg/mL for 5 min at room temperature. The reaction was quenched by the addition of 10 mM phenylmethylsulfonyl fluoride (PMSF). To assess protection by in specific nucleotide states, samples were pre-incubated at 37°C for 45 min with either 1 mM ATP and 5 mM MgCl_2_ (ATP state) or 1 mM ATP, 5 mM MgCl_2_, 5 mM Al(NO_3_)_3_, and 30 mM NaF (ATP-AlFx state) prior to PK addition. Samples were analyzed by 15% SDS-PAGE and Coomassie blue staining. Data analysis was performed using Image J and GraphPad Prism 10.

### NADH-coupled ATPase activity assay

ATP hydrolysis rate were measured using an NADH-coupled regeneration assay^56^. In this system, the regeneration of hydrolyzed ATP by pyruvate kinase is coupled to the oxidation of NADH by lactate dehydrogenase, such that the rate of NADH oxidation is directly proportional to the rate of ATP hydrolysis. Reactions were performed in triplicate with 0.2 μM hTRiC or 0.04 μM yTRiC complex in a buffer containing 20 mM HEPES/NaOH pH 7.5, 50 mM NaCl, 5 mM MgCl_2_, 5% glycerol, and 1 mM ATP. Assays were conducted at 37°C in a 200 μL reaction volume using a 96-well plate reader. The decrease in NADH absorbance at 340 nm was monitored over time, and data were analyzed using GraphPad Prism 10.

### Cell viability assay

Cell proliferation was assessed using the Cell Counting Kit-8 (CCK8; HY-K0301, MCE) according to the manufacturer’s instruction. Cells were seeded at a density of 20,000 cells per well in 100 μL of DMEM supplemented with 10% FBS in 96-well plate (Day 0). At indicated time points, 10 μL of CCK-8 reagent was added to each well (96-well plate), and the plates were incubated at 37 °C for 3.5 h. Absorbance at 450 nm was measured using a Synergy Neo reader.

### Label-free quantification proteomics of HEK293F cell

Cell lysis and protein digestion were performed using the filter-aided sample preparation (FASP) method^57^ with minor modifications. In brief, total protein from each group was diluted in 200 μL of UA buffer (8 M urea, 50 mM Tris−HCl, pH 8.0), loaded onto a 10 kDa ultrafiltration centrifuge tube (Sartorius), and centrifuged at 14,000 ×g for 30 min. This step was repeated twice. Proteins were then alkylated by adding 100 μL of UA buffer containing 10 mM Tris (2-carboxyethyl) phosphine (TCEP) and 20 mM 2-chloroacetamide (CAA), followed by incubation at room temperature with continuous shaking (650 rpm) for 30 min. The buffer was exchanged by adding 300 μL of UA buffer and centrifuged at 14,000 ×g for 30 min (repeated twice), followed by three washes with 300 μL of 50 mM NH_4_HCO_3_ (centrifugation at 14,000 ×g for 20 min). For digestion, 2 μL of trypsin solution (1 μg trypsin in 100 μL of 100 mM NH_4_HCO_3_) was added, and the samples were incubated at 37 °C with shaking (650 rpm) for 12 h. Peptides were eluted by centrifugation at 14,000 ×g for 20 min into a new collection tube. The filtrate was acidified with 0.1% trifluoroacetic acid (TFA), desalted using C18 cartridges (Thermo Scientific) and the concentration was determined using the Pierce Quantitative Colorimetric Peptide Assay kit (Thermo Scientific).

Peptides were analyzed on a hybrid trapped-ion mobility spectrometry (TIMS) quadrupole timeof-flight mass spectrometer (MS) (TIMS-TOF Pro 2, Bruker Daltonics) using a CaptiveSpray nano-electrospray ion source. Samples were dissolved in 10 μL of 0.1% (v/v) formic acid and loaded onto a pulled emitter column (25 cm × 75 μm, 1.7 μm, C18; IonOptiks) maintained at 50 °C. Mobile phases consisted of 0.1% (v/v) formic acid in water (phase A) and acetonitrile (phase B). Peptides were separated using a 60-min stepped gradient at a flow rate of 300 nL/min: 2 to 22% B over 45 min, 22 to 37% B over 5 min, and 37 to 80% B over 5 min. The TIMS-TOF Pro 2 instrument was operated in data-independent acquisition (DIA)-parallel cumulative serial fragmentation (PASEF) mode containing 32 isolation windows (14 m/z width, including a margin of 0.5 m/z). The PASEF method was defined with a range of mobility values of 0.6−1.6 1/K0; a TIMS accumulation time and ion mobility separation were both set at 50 ms. The MS and MS/MS spectra were recorded from m/z 363.9 to 1161.9 Da.

Raw mass spectrometrydata were processed using Spectronaut 19 software. Peptide sequences (and hence protein identity) were determined by searching against the human protein database from the uniport website (https://www.uniprot.org/). Enzyme specificity was set to fully tryptic and semitryptic N-ragged (cleavage C-terminus to Lys and Arg) parameters with a maximum of two missed cleavages permitted. Carbamidomethyl of cysteine (C) was set as a fixed modification, while oxidation of methionine (M) was set as a variable modification. Interference correction was enabled to remove interfering signals, with a minimum of three peptides required for quantitation. Deisotoping was performed based on the RT apex distance and m/z spacing without demultiplexing. Each observed fragment ion was assigned to a single precursor peak. A false discovery rate FDR of 1% was applied based on a target decoy approach.

The Volcano plot is generated by VolcaNoseR website^58^, the Gene Ontology (GO) analysis and protein-protein interaction (PPI) networks were generated in STRING^59,60^.

### Statistics and Reproducibility

Cryo-EM data collection and processing statistics are summarized in Table. S1. Statistical analysis and graphing were performed using GraphPad Prism 10 software, and band densitometry was quantified using ImageJ. For HEK293F growth curves, OD values were analyzed using a two-way ANOVA. For Protease K protection and ATP hydrolysis assays, statistical significance was determined using a one-way ANOVA. All biochemical assays were performed as three independent experiments.

For quantitative proteomics, protein abundance in CCT2-T400P cells was compared to that in WT cells. Significance was evaluated using a two-tailed Student’s t-test based on three independent biological replicates (n=3).

Quantitative data are presented as mean ± standard deviation. Statistical significance is indicated as follows: ns (not significant), p > 0.05; *, p < 0.05; **, p < 0.01; ***, p < 0.001; ****, p < 0.0001.

## Data availability

All data presented in this study are available within figures and in Supplementary Information. Cryo-EM maps and models have been deposited at the Electron Microscopy Data Bank and Protein Data bank with accession codes EMDB-****/PDB-*** for****, EMDB-****/PDB-*** for****, EMDB-****/PDB-*** for****, EMDB-****/PDB-*** for****, and EMDB-****/PDB-*** for****. Detailed information is refered to the Table S1.

## Acknowledgements

We thank the staff of the Electron Microscopy, Database and Computing, Protein Expression and Purification, and the Integrated Laser Microscopy System facilities at the National Facility for Protein Science in Shanghai (https://cstr.cn/31129.02.NFPS) for their instrument support and technical assistance. Special thanks to Prof. Jinsong Li lab and Dr. Diao Lei from Prof. Bao Lan lab for providing the CRISPR/Cas9 vector and offering valuable guidance on cell line construction. This work was supported by grants from Strategic Priority Research Program of CAS (XDB0570303), National Key R&D Program of China (2024YFA1306204 and 2024YFA1803102), the NSFC (32130056 and 32570924), and Shanghai Pilot Program for Basic Research from CAS (JCYJ-SHFY-2022-008).

## Author contributions

C.L., Y.C., and Q.-Y.Z. conceived the project. Q.-Y.Z. generated the CCT2-T400P HEK293F cell line with assistance from Y.-K.W.. C.L. led the construction of the CCT2-T394P-R510H yeast strain, with contributions from Q.-Y.Z., Y.-X.W., and X.Y. Q.-Y.Z. purified the WT and mutant hTRiC, with assistance from W.J. and Q.S. C.L. purified the WT and mutant yTRiC. Q.-Y.Z. prepared hTRiC cryo-EM sample and collected data. C.L. prepared yTRiC cryo-EM samples, collected data, and performed initial data processing. Q.-Y.Z. performed the structure reconstruction, model building, and data analysis. Q.-Y.Z. conducted biochemistry analyses with involvement of Q.Z. Proteomic profiling was carried out by Q.-Y.Z. and S.W., with X.Z. assisting in data analysis. Q.-Y.Z. and Y.C. wrote the manuscript wih input from C.L..

## Competing interests

All authors declare no competing interests.

## Figures

**Fig. S1.**
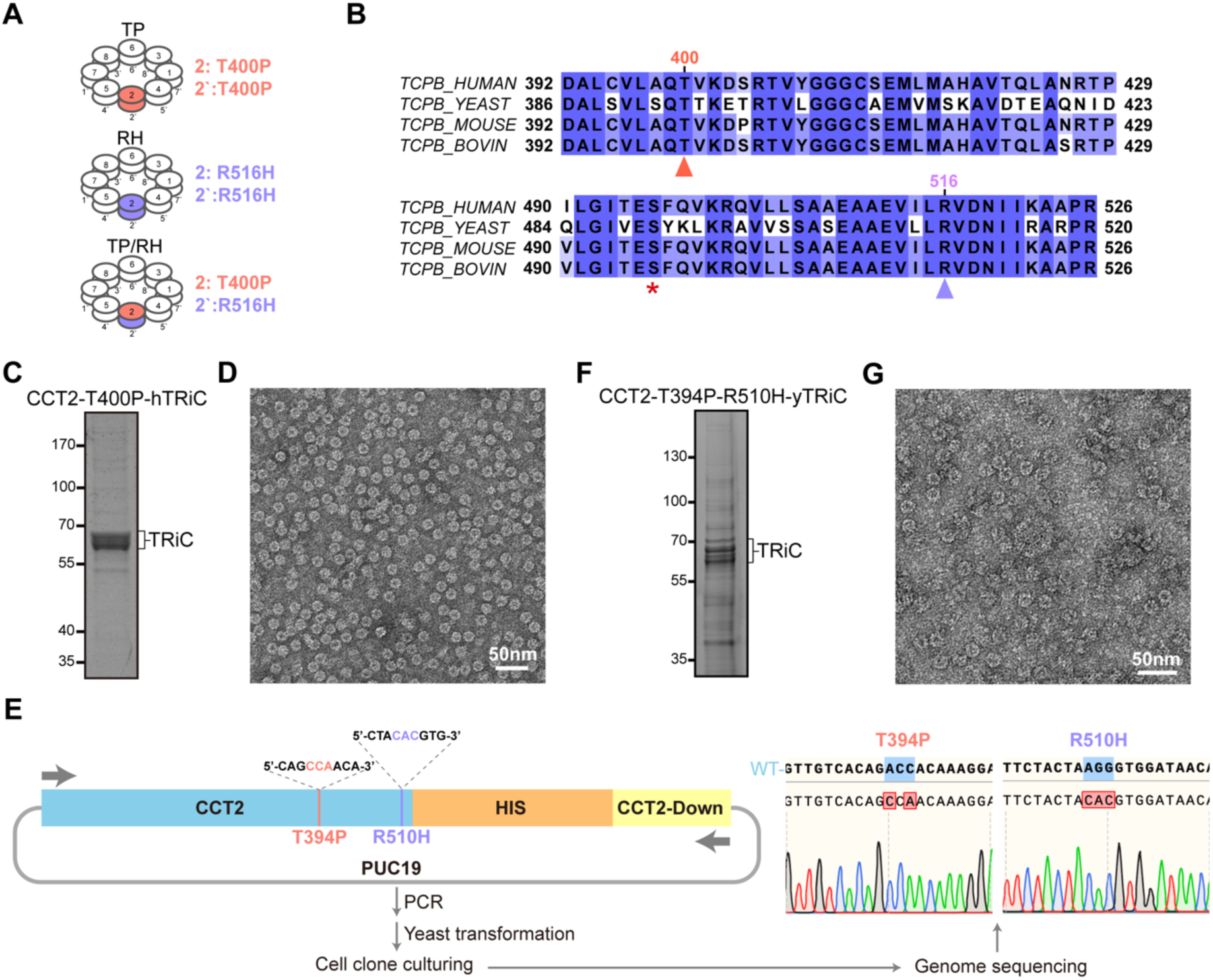
Mutant TRiC complexes purified from HEK293F cells and yeast. (A) Schematic representation of the TRiC variants present in patients with LCA (due to compound heterozygous mutations). (B) Sequence alignment of CCT2 across different species surrounding residues T400 (indicated in red) and R516 (indicated in medium purple); the conserved S496 residue is also labeled (indicated by star). (C) Coomassie blue stained SDS-PAGE of the purified CCT2-T400P hTRiC complex. (D) Negative-stain EM image of the purified CCT2-T400P hTRiC. (E) Workflow for generating the CCT2-T394P-R510H mutant yeast strain via homologous recombination, including sequencing validation of the successfully engineered locus. (F) Coomassie blue-stained SDS-PAGE of the purified CCT2-T394P-R510H yTRiC. (G) Negative-stain EM image of purified CCT2-T394P-R510H yTRiC.

**Fig. S2.**
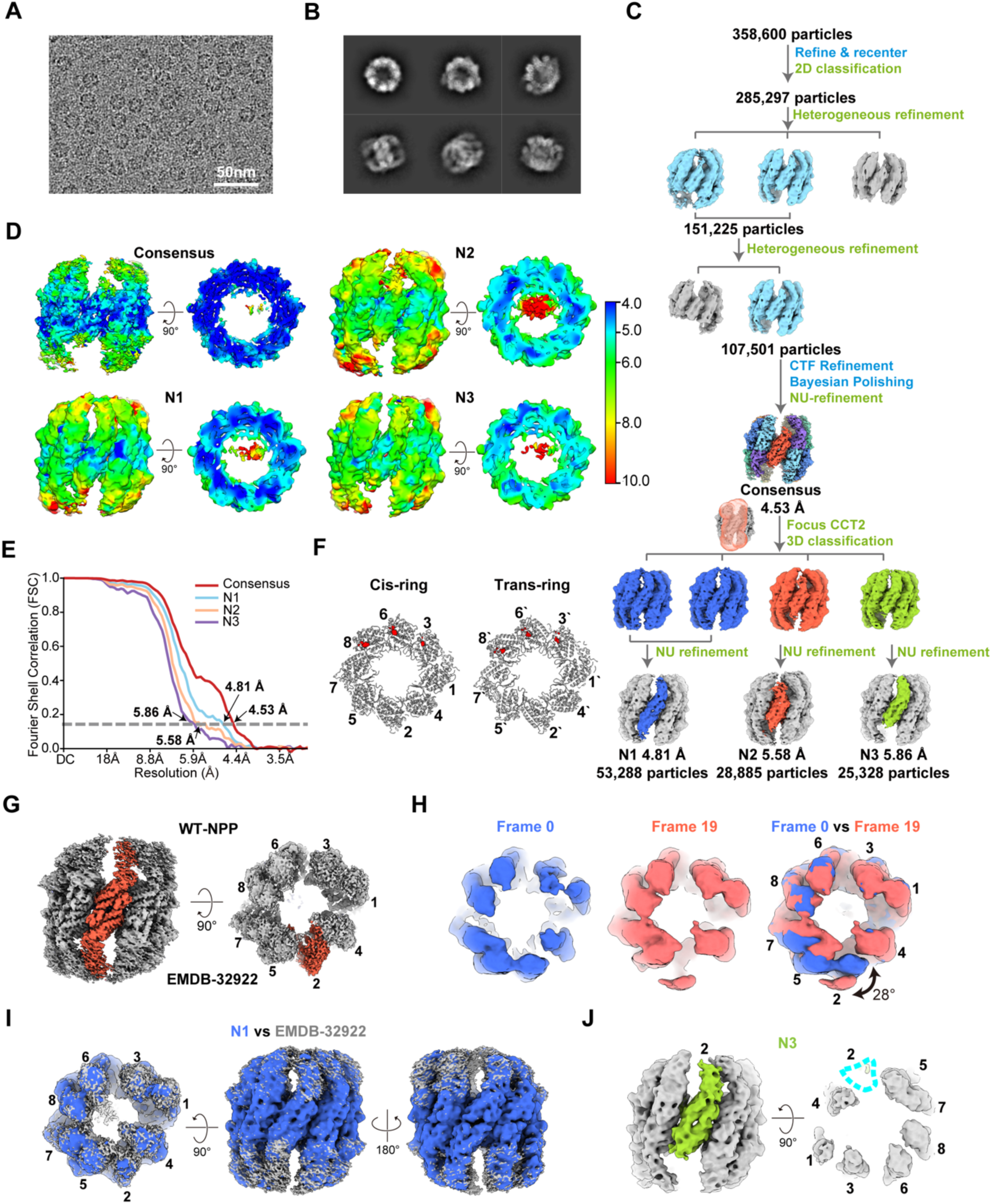
Cryo-EM processing of the CCT2-T400P hTRiC complexes in the NPP state. (A) Representative cryo-EM micrograph of CCT2-T400P hTRiC in the NPP state. (B) Reference-free 2D class averages. (C) Data processing workflow for the CCT2-T400P hTRiC NPP state dataset. (D) Local resolution estimation for the consensus map and the classified N1, N2, and N3 states, displayed as side (left) and central slice top views (right). (E) FSC curves for the final reconstructions. (F) Nucleotide occupancy analysis showing clear nucleotide densities (red) in the CCT6 and CCT8 pockets, with weaker density in CCT3. (G) WT hTRiC in the NPP state (EMDB: 32922), highlighting the resolved CCT2 A/I-domain densities for comparison. (H) 3DVA analysis demonstrating the movement of the mutant CCT2 A-/I-domains. (I) Overlay of N1 state map (blue) and the WT NPP state hTRiC map (gray, EMDB: 32922), indicating no obvious conformational differences between the two structures. (J) Side and end-on views of N3 state, illustrating the missing A- and I-domains in the trans-ring CCT2.

**Fig. S3.**
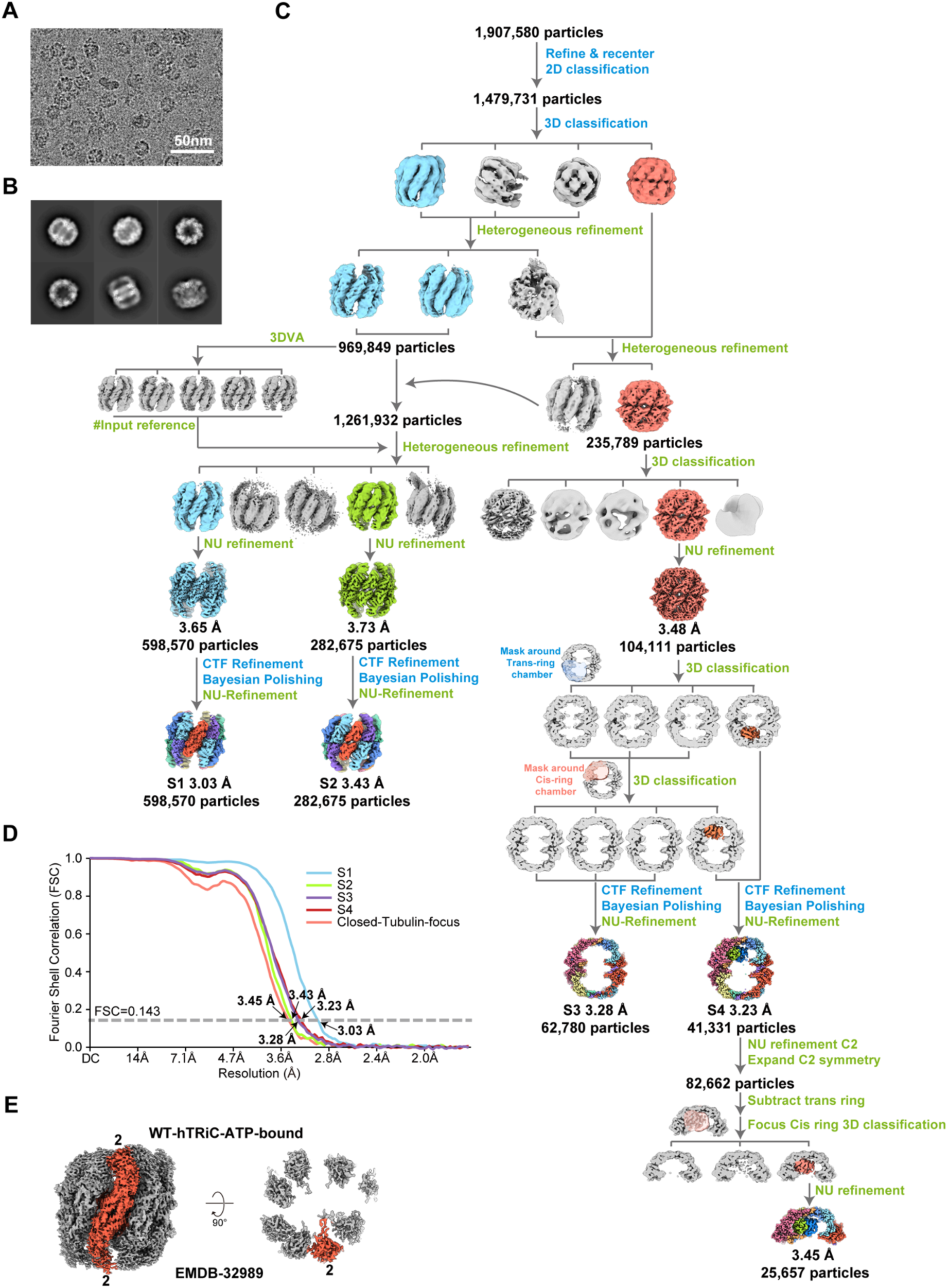
CCT2-T400P hTRiC complexes in the presence of ATP-AlFx. (A) Representative cryo-EM micrograph of CCT2-T400P hTRiC in the presense of ATP-AlFx. (B) Reference-free 2D class averages. (C) Data processing workflow for the CCT2-T400P hTRiC in the presense of ATP-AlFx dataset. (D) FSC curves of the CCT2-T400P hTRiC ATP-AlFx reconstructions. (E) WT hTRiC in the ATP-bound state (EMDB: 32989) showing stabilized densities for the CCT2 A/I-domain.

**Fig. S4.**
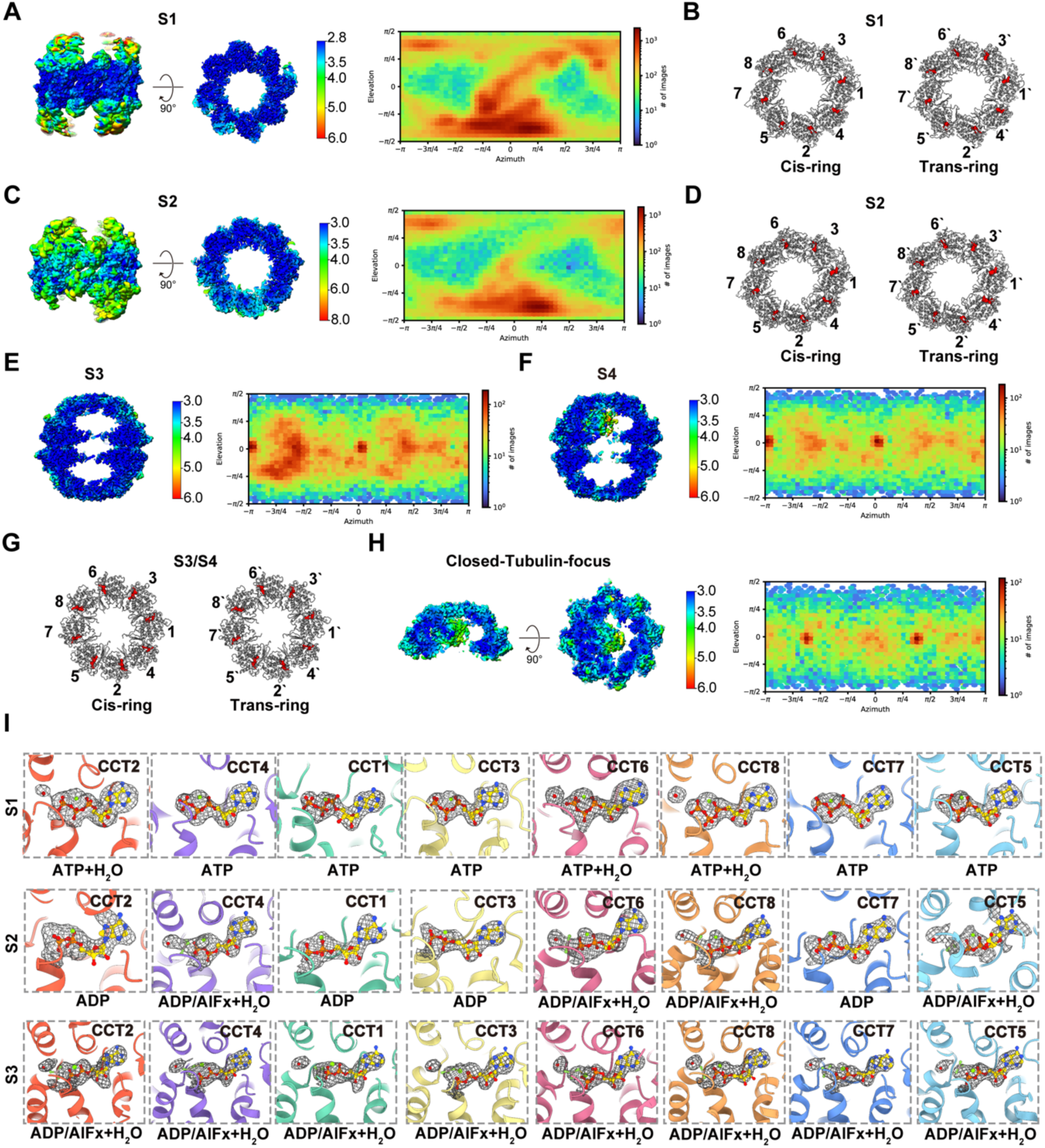
Cryo-EM map quality and structural analysis of CCT2-T400P hTRiC in the presense of ATP-AlFx. (A) Local resolution estimation of the CCT2-T400P TRiC in the S1 state, shown as side (left) and central slice top views (right), along with particle angular distribution. (B) Nucleotide occupancy analysis of the S1 state, with nucleotide densities (red) observed in the pockets of all subunits. (C) Local resolution esitimation and angular distribution of the S2 state. (D) Nucleotide occupancy in the S2 state, with nucelotides bound in the pockets of all subunits. (E-F) Local resolution estimation and particle angular distribution of CCT2-T400P hTRiC in the S3 and S4 state, featuring an empty chamber in (E) and an α-tubulin-occupied chamber in (F). (G) Nucleotide occupancy analysis of the S3 and S4 states, with nucleotide densities observed in the pockets of all subunits. (H) Local resolution estimation and particle angular distribution of the α-tubulin-occupied half-ring density. (I) Nucleotide pocket analysis in the S1 state reveals ATP and a magnesium ion (green sphere) bound in all subunits, with attacking water molecules (red sphere) in CCT2, CCT6, and CCT8. In the S2 state, CCT2/1/3/7 bind ADP, while other subunits bind ADP-AlFx, a magnesium ion, and an attacking water molecule. In the S3 state, all subunits bind ADP-AlFx, a magnesium ion, and an attacking water molecule.

**Fig. S5.**
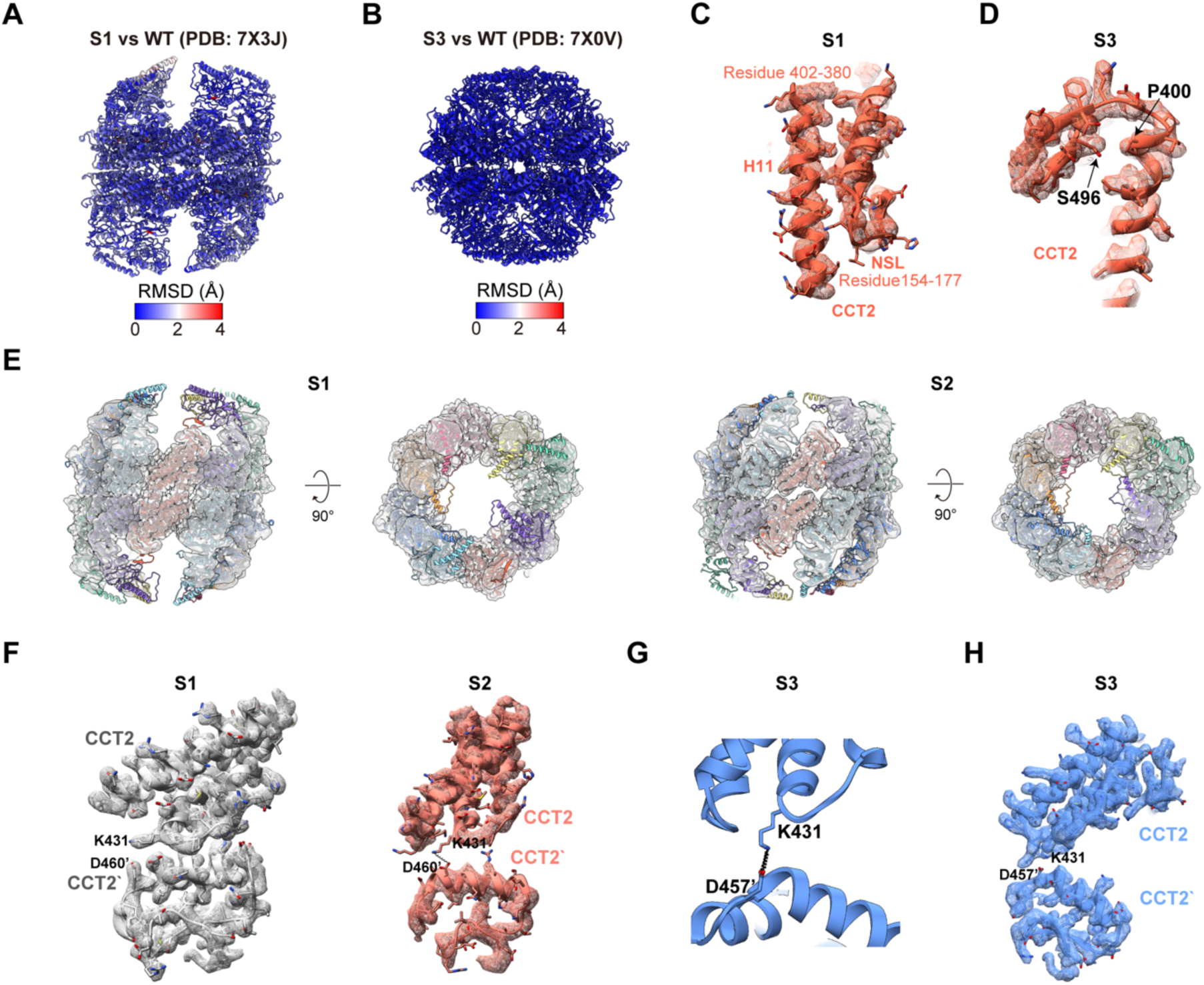
Characterization of the newly identified intermediate state. (A) RMSD analysis comparing the open S1 state of CCT2-T400P hTRiC to WT hTRiC in the open state (PDB: 7X3J). (B) RMSD analysis comparing the closed S3 state of CCT-T400P hTRiC to WT hTRiC in the closed state (PDB: 7X0V). (C-D) Model-map fitting around residue P400 and H11 in the S1 (C) and S3 (D) states. (E) Model-map fitting of the S1 and S2 states, viewed from the CCT2 side and end-on views. (F) Model-map fitting of the CCT2-CCT2’ interaction in the S1 and S2 states. (G) Salt bridge interactions within the CCT2 pair in the S3 state, highlighting the shift to the inner cavity K431-D457’ pair. This illustrates a progressive reorganization of the interaction network from the S1 to S3 states, relative to the configurations shown in Fig. 2N-O. (H) Model-map fitting of the CCT2-CCT2’ interaction in S3 state.

**Fig. S6.**
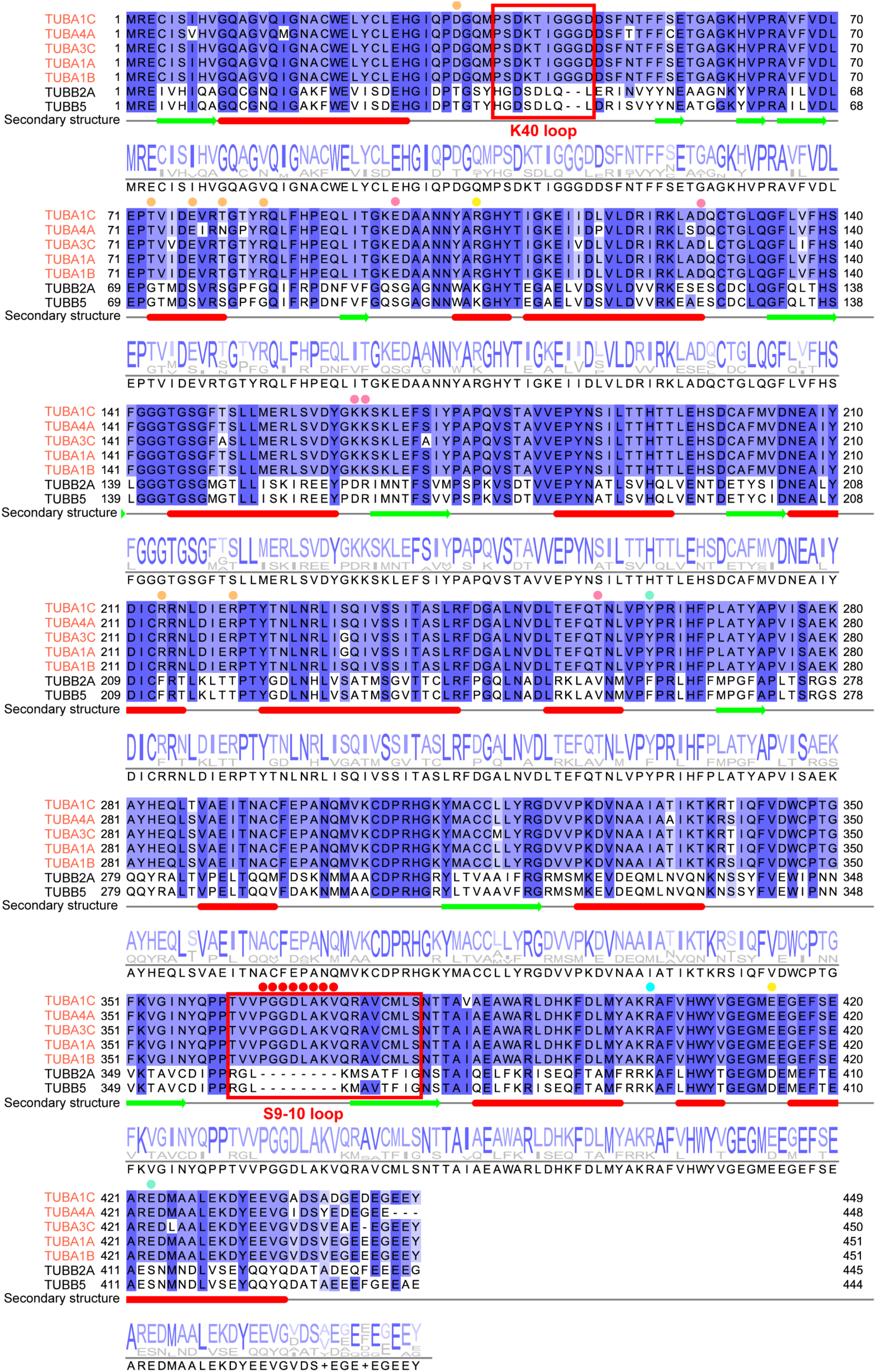
Sequence alignment of α- and β-tubulin subtypes. The sequence alignment highlights the differences between α- and β-tubulin, particularly the longer K40 loop and S9-10 loop, which are unique to α-tubulin. Additionally, it underscores the specifity of the CCT8 interaction sites on α-tubulin, involving residues R214 and R221. Interaction sites on TUBA1C that are unique to α-tubulin are labeled in different color according to the interacting TRiC subunit: orange for CCT8, pink for CCT6, yellow for CCT3, and aquamarine for CCT1. The alignment also reveals high sequence identity between the α-tubulin subtypes TUBA1C and TUBA4A. Since TUBA1C showed higher abundance in the mass spectrometry data, it was selected for building the atomic model.

**Fig. S7.**
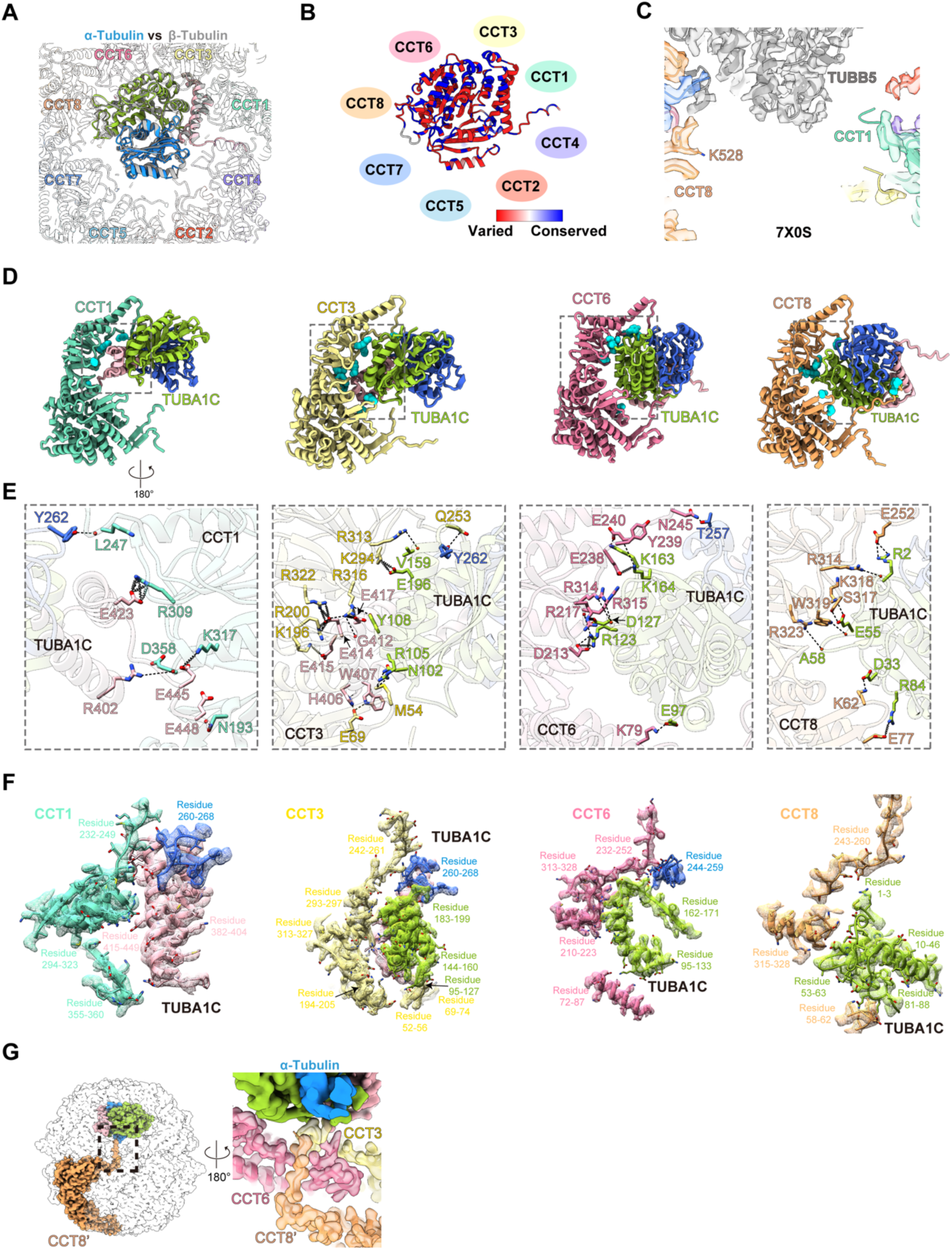
Features of the α-tubulin within the CCT2-T400P hTRiC chamber. (A) Superimposition of α- and β-tubulin within the TRiC chamber reveals their consistent placement on the CCT3/6/8 side (β-tubulin model from PDB: 7X0S). (B) Illustration of α-tubulin’s position in the TRiC chamber, colored by sequence conservation between TUBA1C and TUBB5. Highly conserved regions are observed at the CCT3/6/8 contact sites. (C) Model-map fitting of TRiC-β-tubulin (PDB:7X0S), showing the absence of CCT8 C-terminal tail density around β-tubulin, in contrast to the interaction seen with α-tubulin (related to Fig. 3F). (D) Ribbon diagrams of the interactions between CCT1/3/6/8 subunits and α-tubulin. TRiC amino acid residues forming H-bonds and salt bridges with α-tubulin are represented as cyan spheres. (E) Magnified views of the boxed regions in (D), highlighting H-bonds and salt bridges between α-tubulin and CCT1/3/6/8. (F) Model-map fitting at the interaction sites between CCT1/3/6/8 and α-tubulin. (G) Interactions of the CCT8’ N-terminus from the opposite ring with CCT3/6 and α-tubulin.

**Fig. S8.**
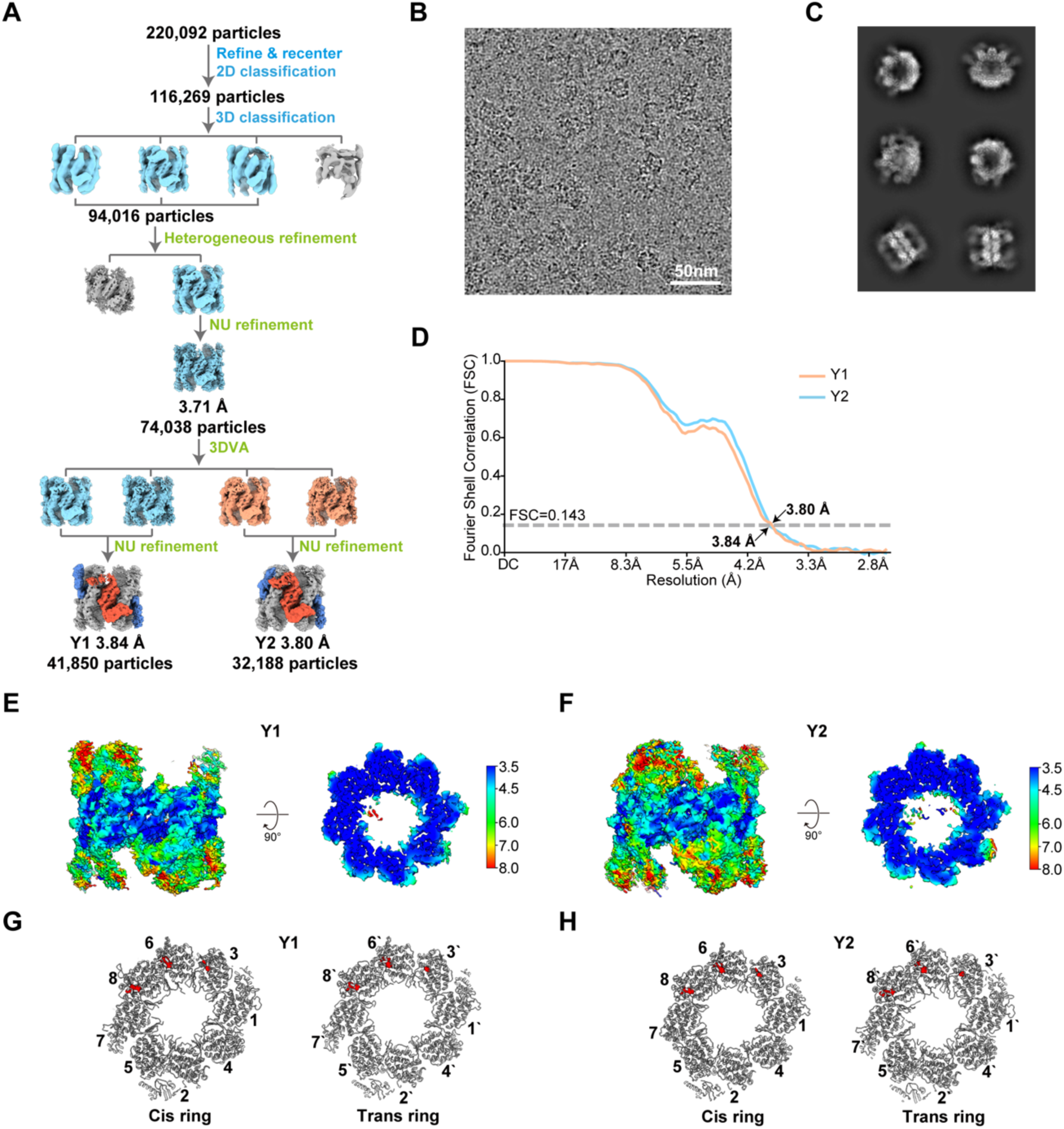
Cryo-EM analysis of CCT2-T394P-R510H yTRiC in the NPP state. (A) Workflow for processing the CCT2-T394P-R510H yTRiC dataset in the NPP state. (B) Representative cryo-EM image. (C) Reference-free 2D class averages. (D) FSC curve of the CCT2-T394P-R510H yTRiC structures in the NPP state. (E-F) Local resolution estimations of the CCT2-T394P-R510H yTRiC for the Y1 (E) and Y2 (F) states, shown as side views (left) and central slice top views (right). (G-H) Nucleotide occupancy in the Y1 (G) and Y2 (H) states, with nucleotide densities (red) observed in the nucleotide pockets of CCT3, CCT6 and CCT8.

**Fig. S9.**
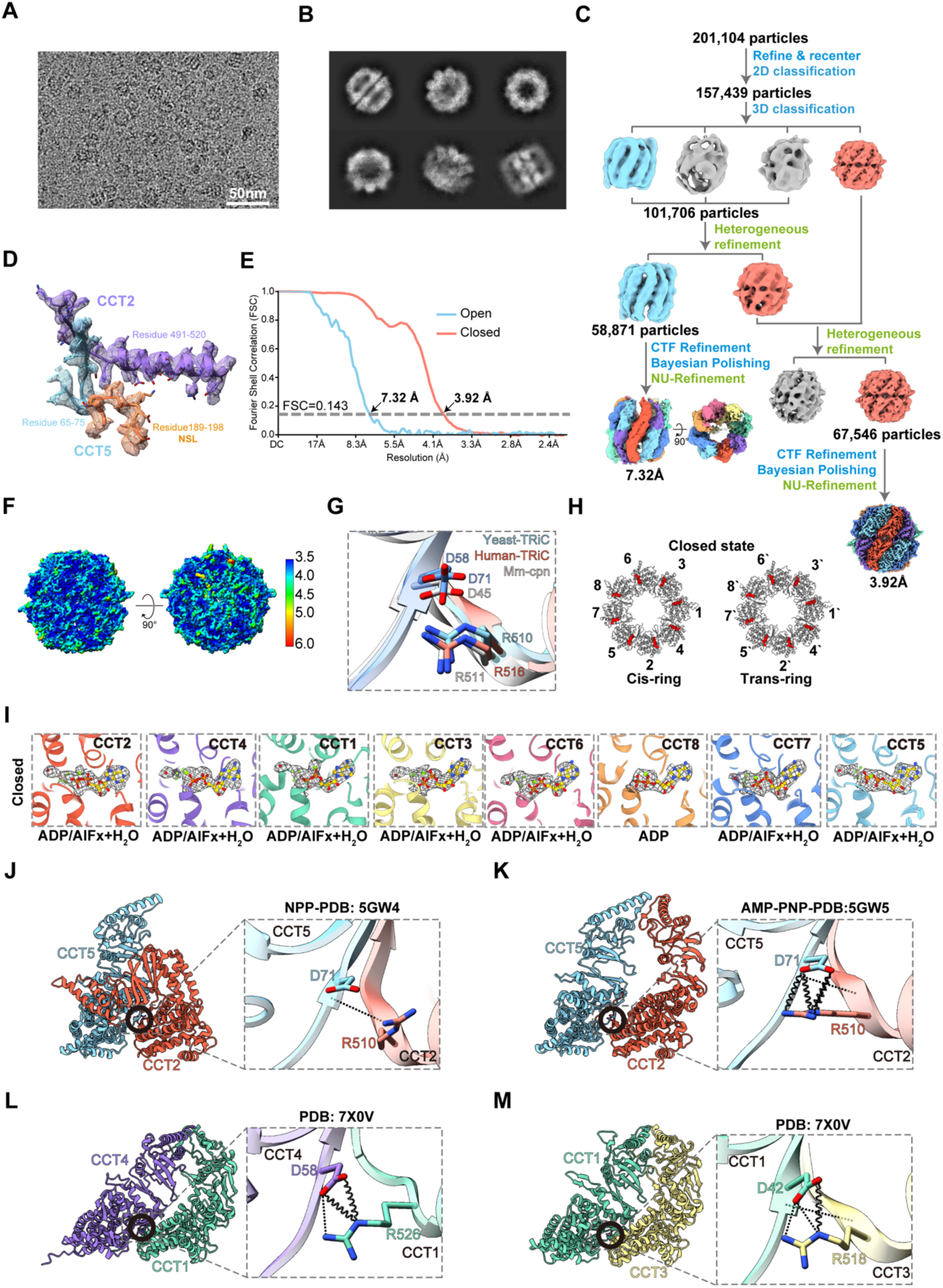
Cryo-EM analysis of CCT2-T394P-R510H yTRiC in the presense of ATP-AlFx. (A) Representative cryo-EM micrograph of CCT2-T394P-R510H yTRiC in the presense of ATP-AlFx. (B) 2D class averages. (C) Workflow for processing the CCT2-T394P-R510H yTRiC dataset in the presense of ATP-AlFx. (D) Model-map fitting of the interaction site between CCT2-H510 and CCT5-D71. (E) FSC curve of the CCT2-T394P-R510H yTRiC structures in the presense of ATP-AlFx. (F) Local resolution estimations of the CCT2-T394P-R510H yTRiC in the closed state, shown as side views (left) and top views (right). (G) The conserved R-D interaction in the CCT2-CCT5 interaction. (H) Nucleotide occupancy in the closed state, with nucelotides bound in the pockets of all subunits. (I) Model-map fitting of the nucleotides: CCT8 binds ADP, while other subunits bind ADP-AlFx, a magnesium ion (green sphere), and an attacking water molecule (red sphere). (J) R-D interaction in CCT2-CCT5 in the NPP state (PDB: 5GW4). (K) R-D interaction in CCT2-CCT5 in the AMP-PNP-bound state (PDB: 5GW5). (L) R-D interaction in CCT1-CCT4 of hTRiC (PDB: 7X0V). (M) R-D interaction in CCT3-CCT1 of hTRiC (PDB: 7X0V).

**Table S1.**
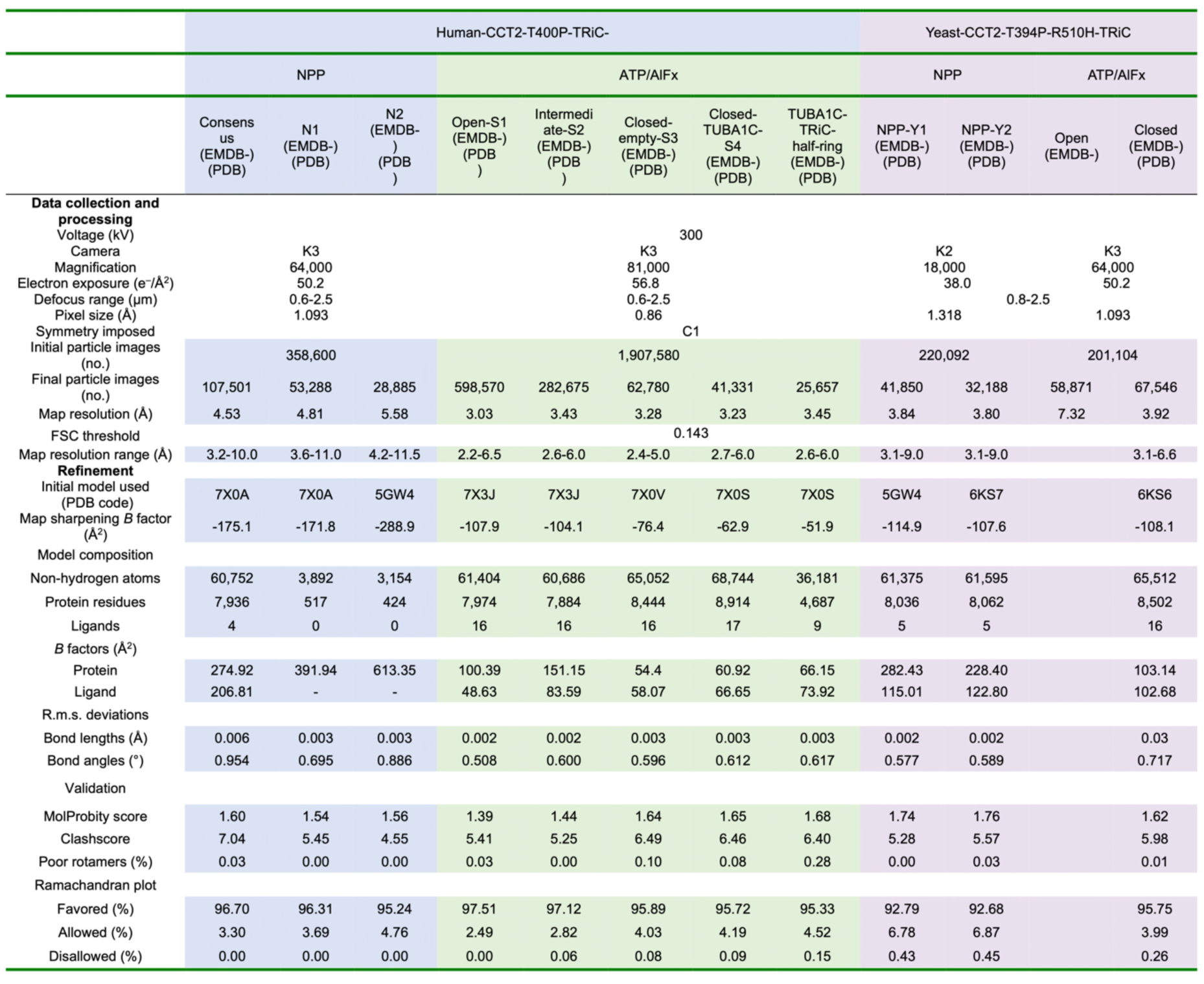
Cryo-EM data collection, processing, and model validation statistics.

**Table S2.** Results of mass spectroscopy (MS) analysis of endogenously purified CCT2-T400P human TRiC with associated tubulin.

|  | Protein | Coverage | PSMs | Peptides | Unique Peptides |
| --- | --- | --- | --- | --- | --- |
| TRiC Subunits | CCT1 | 65 | 943 | 33 | 33 |
|  | CCT2 | 70 | 1243 | 40 | 40 |
|  | CCT3 | 74 | 1278 | 50 | 5 |
|  | CCT4 | 61 | 743 | 37 | 36 |
|  | CCT5 | 66 | 1433 | 42 | 41 |
|  | CCT6 | 61 | 939 | 33 | 26 |
|  | CCT7 | 61 | 984 | 37 | 37 |
|  | CCT8 | 83 | 1203 | 45 | 45 |
| α-Tubulin | TUBA1C | 44 | 100 | 19 | 8 |
|  | TUBA4A | 41 | 91 | 15 | 4 |
| β-Tubulin | TUBB2C | 67 | 182 | 23 | 1 |
|  | TUBB2B | 66 | 180 | 22 | 1 |
|  | TUBB2A | 66 | 179 | 22 | 1 |
|  | TUBB4A | 59 | 151 | 20 | 2 |
|  | TUBB6 | 32 | 85 | 14 | 6 |
| γ-Tubulin | TUBG1 | 20 | 12 | 7 | 7 |

**Table S3.** Interactions between TUBA1C and CCT1/3/6/8 subunits analyzed by PISA.

|  | TRiC subunits |  | TUBA1C |  | Interaction | Distance (Å) |
| --- | --- | --- | --- | --- | --- | --- |
|  | Residue | Atom | Residue | Atom |  |  |
| CCT1 | ARG 309 | [NH2] | GLU 423 | [OE1] | H-bond | 2.49 |
|  | ASN 193 | [ND2] | GLU 448 | [O] | H-bond | 3.54 |
|  | LYS 247 | [O] | TYR 262 | [OH] | H-bond | 3.78 |
|  | ASP 358 | [O] | ARG 402 | [NH2] | H-bond | 3.36 |
|  | ARG 309 | [NE] | GLU 423 | [OE1] | Salt bridge | 3.78 |
|  | ARG 309 | [NH1] | GLU 423 | [OE1] | Salt bridge | 3.94 |
|  | ARG 309 | [NE] | GLU 423 | [OE2] | Salt bridge | 3.83 |
|  | ARG 309 | [NH2] | GLU 423 | [OE2] | Salt bridge | 3.72 |
|  | LYS 317 | [NZ] | GLU 445 | [OE1] | Salt bridge | 3.79 |
| CCT3 | ARG 316 | [NE] | TYR 108 | [OH] | H-bond | 3.82 |
|  | ARG 313 | [NH2] | VAL 159 | [O] | H-bond | 2.96 |
|  | GLN 253 | [NE2] | TYR 262 | [OH] | H-bond | 3.90 |
|  | ARG 316 | [NH1] | GLY 412 | [O] | H-bond | 2.79 |
|  | ARG 200 | [NH2] | GLU 414 | [OE1] | H-bond | 2.60 |
|  | ARG 200 | [NE] | GLU 414 | [OE2] | H-bond | 2.84 |
|  | ARG 316 | [NH1] | GLU 414 | [OE2] | H-bond | 3.46 |
|  | MET 54 | [SD] | ASN 102 | [ND2] | H-bond | 2.73 |
|  | MET 54 | [SD] | ARG 105 | [NH1] | H-bond | 2.96 |
|  | GLU 69 | [OE2] | HIS 406 | [NE2] | H-bond | 2.79 |
|  | GLU 69 | [OE2] | TRP 407 | [NE1] | H-bond | 2.93 |
|  | LYS 294 | [NZ] | GLU 196 | [OE1] | Salt bridge | 3.76 |
|  | LYS 294 | [NZ] | GLU 196 | [OE2] | Salt bridge | 3.48 |
|  | ARG 200 | [NE] | GLU 414 | [OE1] | Salt bridge | 3.12 |
|  | ARG 322 | [NH1] | GLU 414 | [OE1] | Salt bridge | 3.09 |
|  | ARG 200 | [NH2] | GLU 414 | [OE2] | Salt bridge | 3.77 |
|  | ARG 322 | [NH1] | GLU 414 | [OE2] | Salt bridge | 3.03 |
|  | ARG 322 | [NH2] | GLU 414 | [OE2] | Salt bridge | 3.22 |
|  | LYS 196 | [NZ] | GLU 415 | [OE1] | Salt bridge | 3.40 |
|  | ARG 316 | [NE] | GLU 417 | [OE2] | Salt bridge | 3.69 |
|  | ARG 316 | [NH1] | GLU 417 | [OE2] | Salt bridge | 3.59 |
|  | Residue | Atom | Residue | Atom |  |  |
| CCT6 | LYS 79 | [NZ] | GLU 97 | [OE1] | H-bond | 3.86 |
|  | ARG 314 | [NE] | ASP 127 | [O] | H-bond | 3.07 |
|  | ARG 314 | [NH2] | ASP 127 | [O] | H-bond | 3.08 |
|  | ASN 245 | [ND2] | THR 257 | [OG1] | H-bond | 2.81 |
|  | ASP 213 | [OD1] | ARG 123 | [NH1] | H-bond | 3.37 |
|  | ASP 213 | [OD2] | ARG 123 | [NH2] | H-bond | 2.47 |
|  | TYR 239 | [O] | LYS 164 | [NZ] | H-bond | 3.62 |
|  | GLU 240 | [OE1] | LYS 163 | [NZ] | H-bond | 2.54 |
|  | ARG 315 | [O] | ARG 123 | [NH1] | H-bond | 2.69 |
|  | ARG 217 | [NE] | ASP 127 | [OD1] | Salt bridge | 3.91 |
|  | ASP 213 | [OD1] | ARG 123 | [NH2] | Salt bridge | 3.52 |
|  | ASP 213 | [OD2] | ARG 123 | [NH1] | Salt bridge | 3.08 |
|  | GLU 238 | [OE1] | LYS 164 | [NZ] | Salt bridge | 3.69 |
|  | GLU 238 | [OE2] | LYS 164 | [NZ] | Salt bridge | 3.47 |
| CCT8 | ARG 314 | [NH2] | GLY 43 | [O] | H-bond | 3.53 |
|  | SER 317 | [OG] | GLU 55 | [OE2] | H-bond | 2.83 |
|  | LYS 318 | [N] | GLU 55 | [OE1] | H-bond | 3.81 |
|  | TRP 319 | [NE1] | GLU 55 | [O] | H-bond | 2.79 |
|  | ARG 323 | [NH1] | GLY 57 | [O] | H-bond | 3.65 |
|  | GLU 252 | [OE2] | ARG 2 | [NH1] | H-bond | 3.66 |
|  | GLU 252 | [OE2] | ARG 2 | [NH2] | H-bond | 3.49 |
|  | ASP 544 | [OD1] | TYR 210 | [OH] | H-bond | 3.39 |
|  | ASP 544 | [OD1] | ARG 214 | [NH1] | H-bond | 3.38 |
|  | ASP 544 | [OD1] | ARG 221 | [NH2] | H-bond | 2.40 |
|  | ASP 545 | [O] | ARG 214 | [NH2] | H-bond | 3.40 |
|  | ASP 545 | [O] | ARG 214 | [NH1] | H-bond | 3.36 |
|  | LYS 62 | [NZ] | ASP 33 | [OD2] | Salt bridge | 3.80 |
|  | GLU 77 | [OE1] | ARG 84 | [NH2] | Salt bridge | 3.79 |
|  | ASP 544 | [OD2] | ARG 221 | [NH2] | Salt bridge | 3.52 |

**Movie S1 Increased CCT2 dynamics caused by the T400P mutation.** A comparison of the N1- and N2-state CCT2 models shows that the major flexibility occurs in helices H6, H7, and H11 of the intermediate domain. As a result, the NSL linking H6 and H7 shifts away from the E-domain.

## Notes

### Competing Interest Statement

The authors have declared no competing interest.

